# The structures of two intermediates *en route* to DNA repair within a (6-4) photolyase

**DOI:** 10.64898/2026.11.04.763790

**Authors:** Chao Yang, Qi Yu, Xiaodan Cao, Sven Falke, Jie Nan, Emma V. Beale, Henry N. Chapman, Dongping Zhong, Thomas J. Lane

## Abstract

Pyrimidine (6-4) pyrimidone photoproducts (6-4PPs) are mutagenic DNA lesions produced when UV light crosslinks two adjacent pyrimidine bases. These lesions must be repaired to maintain genome integrity, a function executed by the (6-4) photolyase enzymes. This repair chemistry is complex, involving the transfer of a hydroxyl group from one base to another, accompanied or followed by cleavage of an interbase bond. It remains unclear how the enzyme orchestrates this reaction. Typically, models invoke a stepwise process in which an oxetane intermediate is transiently populated, but despite over thirty years of investigation, such an intermediate has never been definitively observed. Conducting dose-controlled X-ray crystallography on the (6-4) photolyase from *Vibrio cholerae*, we observed the transformation of 6-4PP into an oxetane species. Furthermore, we identified a previously unanticipated hemi-cleaved intermediate. The progressive transformation of 6-4PP into oxetane, hemi-cleaved, and fully repaired species requires a reexamination of the existing models of DNA repair by (6-4) photolyases.

## Introduction

The ultra-violet (UV) component of sunlight is highly damaging to genetic material^1–4^. Specifically, UV-driven chemistry produces mutagenic DNA lesions, primarily cyclobutane pyrimidine dimers (CPDs) and pyrimidine (6-4) pyrimidone photoproducts (6-4PPs)^3,4^. Whereas a CPD joins adjacent bases through a compact cyclobutane ring, a 6-4PP contains a highly distorted interbase linkage and an altered 3′ pyrimidine (Fig. 1a). Both lesions disrupt base pairing and DNA stacking^5,6^ and can alter replication^7^ and transcription^8^. In many organisms, these lesions are repaired by photolyases, a class of flavoproteins, through a light-dependent process known as photoreactivation^9^. Specifically, these enzymes consume blue light to drive electron transfer from a fully reduced flavin adenine dinucleotide (FADH⁻) to the damaged DNA bases, thereby triggering a chemical transformation of the lesion back to its functional, unmodified state^9^.

**Fig. 1.**
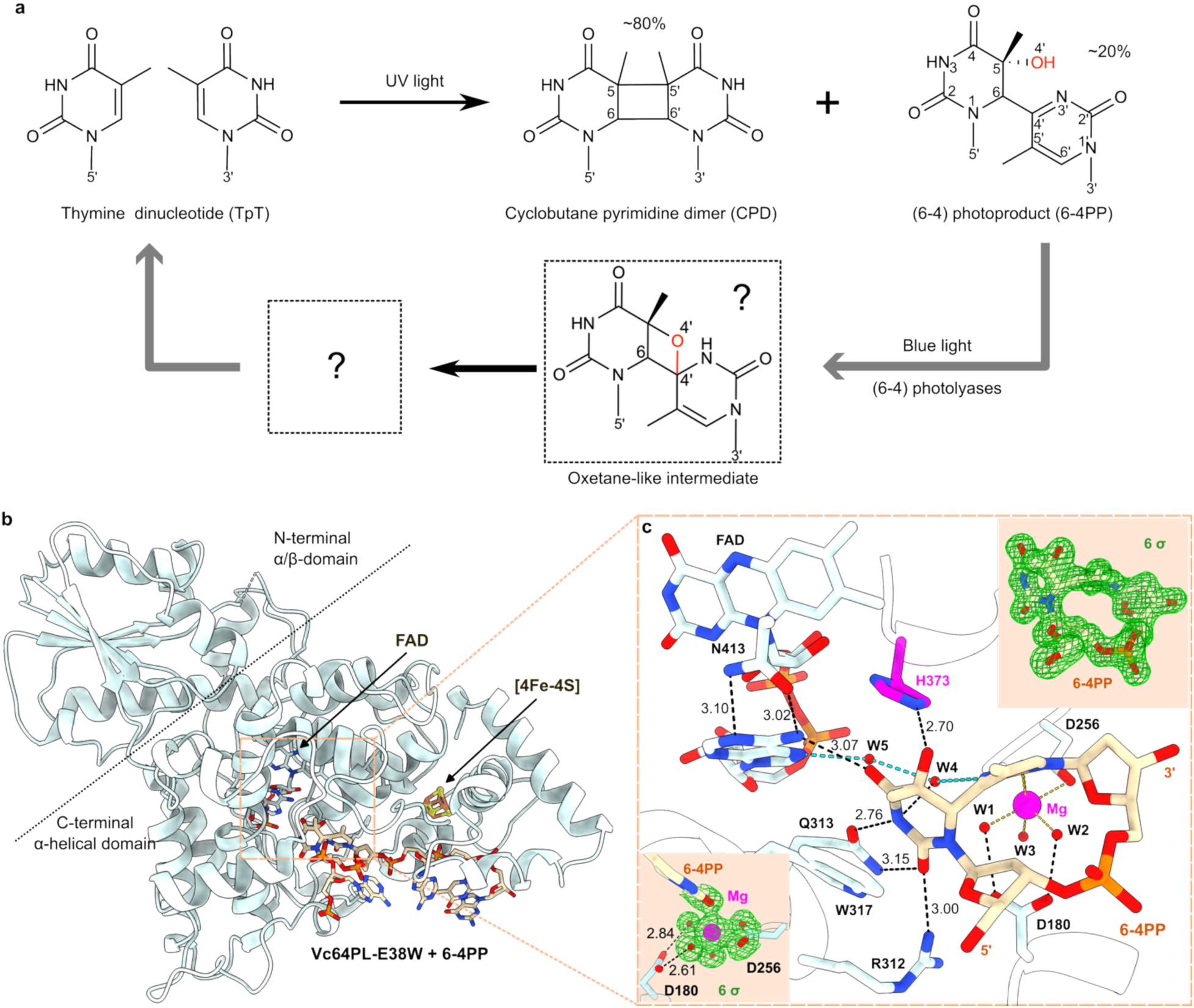
**a, 6-4PP formation and repair.** Exposure to UV light leads to the formation of two common DNA lesions: CPDs and 6-4PPs. (6-4) photolyases can repair 6-4PPs specifically. Many repair models^10^ invoke an oxetane-like species as the key intermediate. **b, Overall structure of Vc64PL-E38W:6-4PP complex.** Structure of Vc64PL-E38W in complex with 6-4PP (PDB code: 31JN) determined using serial femtosecond crystallography and refined to 1.67 Å, showing the overall fold of the protein, the bound flavin adenine dinucleotide (FAD) cofactor, the iron-sulfur cluster ([4Fe-4S]), and the bound DNA that contains a 6-4PP lesion. **c, The active site**, enlarged to show the major interactions between 6-4PP and the active-site residues. Black and cyan dashed lines represent hydrogen-bonding interactions (< 3.5 Å), with distances shown in Å. Cyan dashed lines highlight a two-water (W4, W5) wire connecting the N3′ atom of the 3′ ring to the adenine moiety of FAD. Gold dashed lines highlight the inner-sphere coordination (2.0–2.2 Å) of a hexa-coordinate Mg^2+^. This Mg^2+^ cluster has a near-octahedral geometry in which the Mg^2+^ is coordinated by the O2′ atom of the 3′ ring, two aspartate oxygens (D256) and three water molecules (W1–W3). Insets are polder maps (contoured at 6 σ) calculated with the 6-4PP lesion and the Mg^2+^-coordinated cluster omitted.

Repair of 6-4PPs poses an exceptional chemical challenge for DNA repair enzymes^10,11^. Successful repair requires precise orchestration of photoinduced electron transfer, proton transfer, reversal of the O4′ (Fig. 1a) hydroxyl transfer, and selective bond cleavage while suppressing competing radical pathways^10–12^. This mechanistic complexity has fueled more than three decades of debate regarding the elementary steps of catalysis^10,13^.

A central unresolved question is how (6-4) photolyases reverse the O4′ hydroxyl transfer. One possibility emerged early on when studies identified an unstable oxetane species as the precursor of some UV-induced thymine-thymine adducts including 6-4PPs^14,15^. After the discovery of (6-4) photolyase^13^, Kim *et al.* proposed that the repair proceeds through an oxetane intermediate, which they suggested would form in the dark and only proceed to fully repaired DNA following photoactivation^16^. Although subsequent crystal structures disproved oxetane formation in the dark^17,18^, this oxetane hypothesis became the centerpiece of prevailing mechanistic models and received substantial support from simulations^19–22^, studies of model compounds^23,24^, and substrate analogue experiments^25,26^.

The current corpus of evidence supporting an oxetane intermediate is substantial, but indirect. Recently, a time-resolved UV/IR study showed spectroscopic signatures consistent with oxetane formation in *Xenopus laevis* (6-4) photolyase^27^. Moreover, oxetane formation is coherent with what is known about the chemistry of the (6-4) photolyase active site. Specifically, a conserved histidine is indispensable for catalysis^28^, and has been proposed to promote oxetane formation by proton transfer^12,20^, consistent with EPR/ENDOR spectroscopy^29^, structures^17,18^, isotope effects and ultrafast spectroscopy^12^. In prevailing models^10^, electron transfer forms a 6-4PP radical anion, histidine-mediated proton transfer promotes oxetane formation, and ring opening reverses hydroxyl transfer. Yet, no oxetane intermediate has been definitively observed during 6-4PP repair.

This ambiguity regarding what metastable intermediates – if any – are populated during repair has made a decisive account of the molecular mechanism of (6-4) photolyases impossible. Structural studies in particular, such as X-ray crystallography, could provide a direct means of visualizing catalytic intermediates, but have so far been limited to stable reaction states^17,18,30^ and have not yet been able to capture metastable species populated during repair. While time-resolved serial crystallography has successfully captured catalytic intermediates during CPD repair^31,32^, the much lower quantum yield^12,33–35^ of (6-4) photolyases has so far precluded analogous structural studies.

Seeking a suitable system to employ in time-resolved crystallography experiments, we screened (6-4) photolyase constructs from several species. In crystals (Fig. S1) of an E38W mutant of *Vibrio cholerae* (6-4) photolyase (Vc64PL) bound to 6-4PP, we observed that synchrotron radiation unexpectedly drove rapid lesion repair; lower-dose datasets contained active-site density consistent with an intermediate (Fig. S2). X-ray radiation is known to generate free electrons that can drive reduction chemistry^36^, and X-ray driven DNA repair has previously been observed in a CPD photolyase^37^. Reasoning that X-ray radiation could initiate repair while cryogenic trapping would enrich the population of interesting intermediates, we characterized E38W:6-4PP crystals under controlled X-ray exposures. This approach captured a reaction pathway of 6-4PP repair, including two intermediates at 1.40 Å resolution.

## Results

### XFEL light yields a protein:DNA complex structure free of accumulated X-ray radiation damage

Vc64PL is a prokaryotic (6-4) photolyase containing FAD, a lumazine antenna (6,7-dimethyl-8-ribityllumazine, DLZ) and a [4Fe-4S] cluster^38^. The E38W mutant lacks the DLZ antenna but is otherwise photoactive and structurally similar to the wild-type enzyme (Fig. S3-S6). E38W readily binds to a single-stranded 8-mer of DNA containing a centrally located 6-4PP lesion, and the resulting complex forms both disk-shaped and rod-shaped crystals (Fig. S1) depending on the crystallization precipitant; both crystal forms are suitable for X-ray analysis.

Before studying these crystals as a function of radiation dose, we sought a ground-state structure free from accumulated X-ray radiation damage that could act as a definitive reference. To do so, we determined the structure of the E38W:6-4PP complex using serial femtosecond crystallography (SFX) at SwissFEL (Table S1).

The structure (PDB: 31JN) was refined to 1.67 Å and resolves the intact 6-4PP lesion, the FAD cofactor, the [4Fe-4S] cluster and an ordered active-site solvent network (Fig. 1b-1c). E38W retains the previously reported wild-type fold^39,40^, which we describe briefly. An N-terminal α/β-domain and a C-terminal α-helical domain bind the two redox-active cofactors, FAD and [4Fe-4S]. At the site of the E38W mutant, the loop containing E38W is distorted and unresolved, consistent with loss of hydrogen bonds with DLZ and internal hydrogen bonds^39–43^. Six bases of the kinked 8-mer DNA strand are ordered; the lesion flips into the active site, inserting itself next to FAD, as in other photolyases^17,18,37,44^. The minimum distance between the bound DNA and the [4Fe-4S] cluster is 9 Å (Fig. S7).

Of potential interest for the mechanism of catalysis, the transferred O4′ hydroxyl lies only 2.70 Å from the Nε2 atom of H373 in an orientation compatible with proton transfer (Fig. 1c). In addition, a pair of water molecules (W4, W5) links the lesion to the adenine moiety of FAD, within a broader ordered hydrogen-bonding network in the active site (Fig. S8). Finally, the crystallization buffer contains magnesium, and a hexa-coordinate Mg^2+^ anchors the 3′ ring of the lesion to D256, resolving an outstanding puzzle as to why Mg^2+^ specifically boosts DNA-repair efficiency of (6-4) photolyases from prokaryotes but not those from eukaryotes^45^. Such Mg^2+^-centered clusters are common in other proteins^46^ but unreported in photolyases^17,18,37,39–42,44^ and absent from our DNA-free E38W structure (PDB: 31NN), suggesting that 6-4PP and Mg^2+^ bind collaboratively to prokaryotic (6-4) photolyases.

Having established a structural baseline, we next aimed to study crystals after the effects of X-ray exposure in order to identify and characterize putative intermediates in the 6-4PP repair mechanism.

### Synchrotron radiation drives 6-4PP repair via two sequential metastable intermediates

In our early attempt to resolve the E38W:6-4PP complex structure, we inadvertently obtained a structure (PDB: 31IF, 0.717 MGy) exhibiting fully repaired DNA when employing conventional cryogenic rotational crystallography on a rod-shaped crystal (space group P2_1_2_1_2_1_) under the full transmission of the X-rays (1.2 × 10^8^ photons/second/µm^2^) at the P14 beamline of PETRA III (Table S2). To isolate and systematically characterize the products of X-ray exposure, in subsequent experiments we carefully controlled the crystal size and X-ray dose.

Using what we termed a “continuous-irradiation experiment” (Fig. 2a), we collected diffraction data from disk-shaped crystals (space group P6_4_22) while systematically increasing the transmission of the X-ray beam from 1% to 40%. The high symmetry of the crystals allowed us to slice rotational data into 80° wedges, each of which contains complete dataset and corresponds to a total exposure time of 6-27 seconds. These were processed individually to produce a dose-resolved series. Many of our crystals diffracted beyond 1.4 Å, which allowed unambiguous assignment of chemical species using polder^47^ maps computed against refined models with the lesion omitted. Ordering datasets by their cumulative dose^48^ produced a smooth progression of electron density in the active site, readily interpreted as sequential conversion starting from the initial reactant (6-4PP) to an oxetane-like intermediate (OXE), followed by a novel species we describe as a hemi-cleaved intermediate (HEI), and finally a genuine, repaired thymine pair (TpT) (Fig. 2b).

**Fig. 2.**
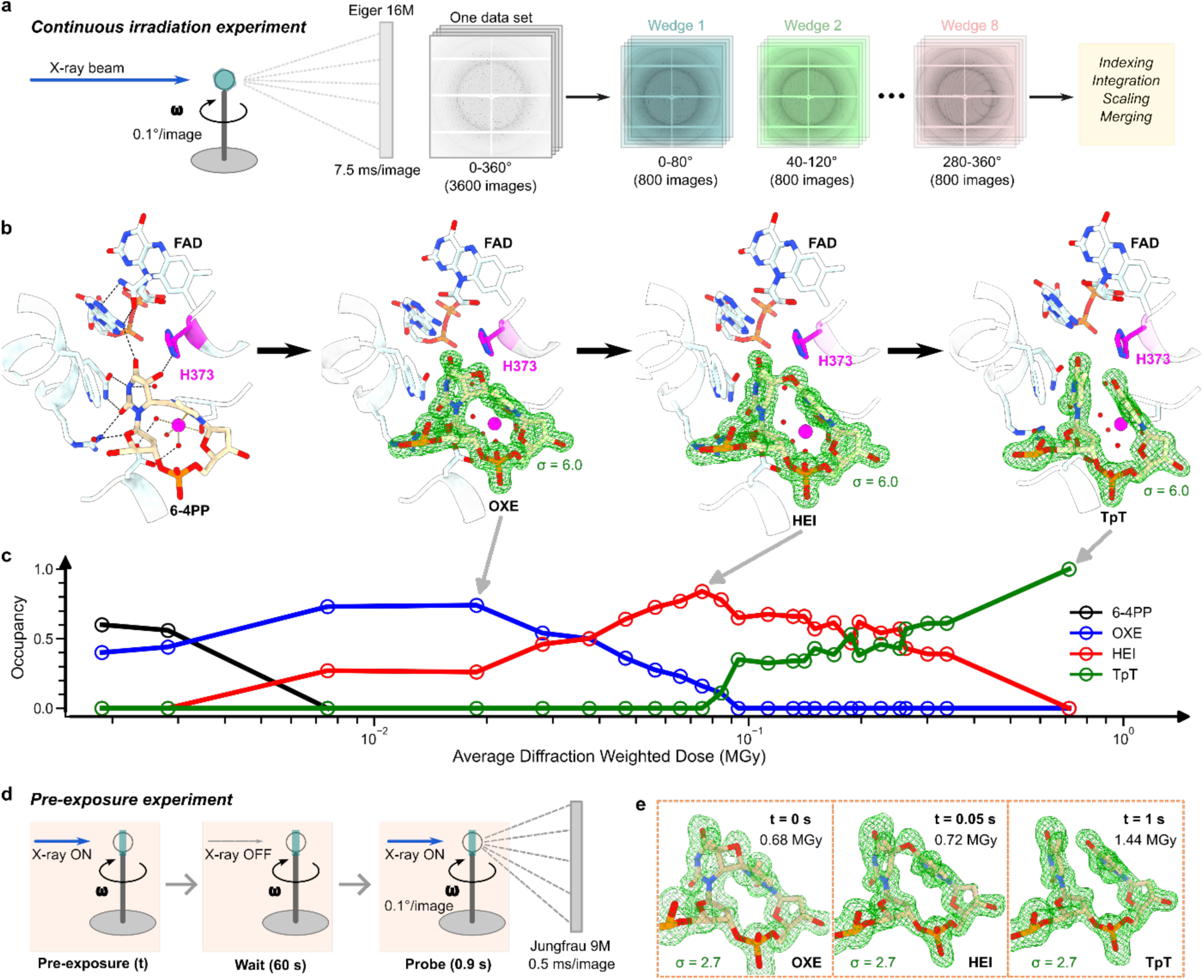
Sequential lesion structures observed in two independent experiments. **(a-c): Continuous-irradiation experiment chronologically orders observed intermediate states. a**, Scheme of experimental setup and dose series data analysis. **b**, Sequential structures show X-ray-induced 6-4PP repair. The SFX structure containing 6-4PP is shown as a reference. An oxetane-like intermediate (OXE, PDB: 31TN), a hemi-cleaved intermediate (HEI, PDB: 31UD) and the repair product (TpT, PDB: 31IF) were captured by gradually increasing X-ray radiation. The maps (green) shown are polder maps contoured at 6 σ, calculated with the lesion model omitted. **c**, Occupancy evolution of 6-4PP, OXE, HEI and TpT as a function of X-ray dose. Here each data point corresponds to a total X-ray exposure time spanning 6 to 27 seconds. **(d-e): Pre-exposure experiment reproduces the sequential lesion structures. d,** Scheme of pre-exposure experiment. **e,** The refined lesion structures and calculated polder maps (contours: 2.7 σ) obtained as the pre-exposure time was increased reveal the OXE, HEI, and TpT species in resolved structures (PDB codes: 32AM, 32AN, 32AO, respectively). For both experiments, dose values reported are average diffraction-weighted (ADW) dose calculated by RADDOSE-3D^48^. For the pre-exposure experiment, the reported doses include radiation from both pre-exposure and data collection (probe).

We built initial coordinate models of OXE and HEI by refining the data wedges (around 0.02 and 0.08 MGy, respectively) that produced electron-density maps that could be unambiguously interpreted as a single species resembling OXE or HEI (Fig. S9-S10). Since most data wedges likely contained mixed states, we used Phenix^49^ to perform occupancy refinement against each data wedge using a multi-state model in which we included the model of the lesion in two different states, and all parameters were frozen except the occupancies of the lesion (see Methods). This refinement against successive wedges showed systematic occupancy changes as dose increased: loss of substrate (6-4PP) accompanied first by accumulation of OXE, followed by HEI and finally TpT (Fig. 2c). This yielded maximum occupancies of 0.74 and 0.84 for OXE and HEI, respectively, in the data wedges corresponding to 0.019 and 0.075 MGy, which were used to obtain our deposited 1.40-Å intermediate structures (PDBs: 31TN, 31UD) containing OXE and HEI.

We speculated that the progression of the repair reaction was not solely determined by the amount of received X-ray dose. Therefore, we performed a “pre-exposure experiment” (Fig. 2d), where X-ray radiation was deposited in a shorter collection time using an intense pink (ΔE/E ∼ 1%) X-ray beam (4.1 × 10^9^ photons/second/µm^2^) at MicroMAX. We executed this experiment on rod-shaped crystals using variable X-ray pre-exposures: no X-ray (control), 0.05 or 1 second, followed by a delay of 60 seconds while the X-rays were shuttered, and finally probed by a 0.9-second diffraction collection covering a 180° rotation. The control experiment produced a 1.50-Å structure (PDB: 32AM, 0.68 MGy) containing OXE; the calculated polder map displayed a clearly resolved central cavity within the four-membered ring (Fig. 2e). With pre-exposures of 0.05 and 1 second, we observed a sequential appearance of HEI and TpT as the total dose was increased to 0.72 and 1.44 MGy, respectively (Fig. 2e, S11-S13, and Table S3).

These two experiments confirmed the sequential progression of OXE, HEI and TpT found in our continuous-irradiation studies. They demonstrated that the observed 6-4PP repair was affected by both X-ray dose and the dose rate, as the 0.68 MGy OXE structure collected in the pre-exposure experiment corresponds to a dose where we observed fully repaired DNA in our continuous-irradiation experiment (Fig. 2c). Further, these experiments show that the X-ray driven OXE formation occurs in less than 1 second and the subsequent transitions from OXE to TpT happen on a timescale of seconds at 100 K.

Having established the sequence of populated intermediates, we next analyzed their structures in detail. While the maps from the pre-exposure experiment are of high quality, ultimately structures refined against the data from the continuous-irradiation experiment produced the highest quality maps and models (Fig. S9 and Table S2). We employed them in our subsequent analysis.

### Observation of an oxetane-like species bound to (6-4) photolyase

Our data revealed unambiguous structural evidence for an oxetane-like species in the active site of E38W (Fig. 3a), which is the first proof of the existence of OXE in a (6-4) photolyase. The calculated polder map supports a model-bias-free assignment of the four-membered oxetane ring, in which an additional C4′-O4′ bond has formed while the pre-existing C5-O4′ bond and interbase C6-C4′ bond are retained (Fig. 3a insert). The intermediate is both well-defined by electron-density signals (Fig. S9 and S10) and present at high occupancy (estimated to be 0.74 from refinement), with the polder map clearly interpretable when contoured to 6 σ. The high quality of this dataset enabled refinement of the oxetane ring geometry. In our OXE structure (PDB: 31TN), the protein backbone, FAD/lesion contacts and [4Fe-4S] cluster remain essentially unchanged when compared to our SFX structure of the E38W:6-4PP complex (Fig. S10 and S14). Of note, the active-site hydrogen-bonding network is unperturbed. Specifically, H373 remains hydrogen-bonded to the transferred O4′ atom, the Mg^2+^ site remains hexa-coordinate in the presence of OXE, and all other interactions between 6-4PP and active-site residues are retained (Fig. 3a).

**Fig. 3.**
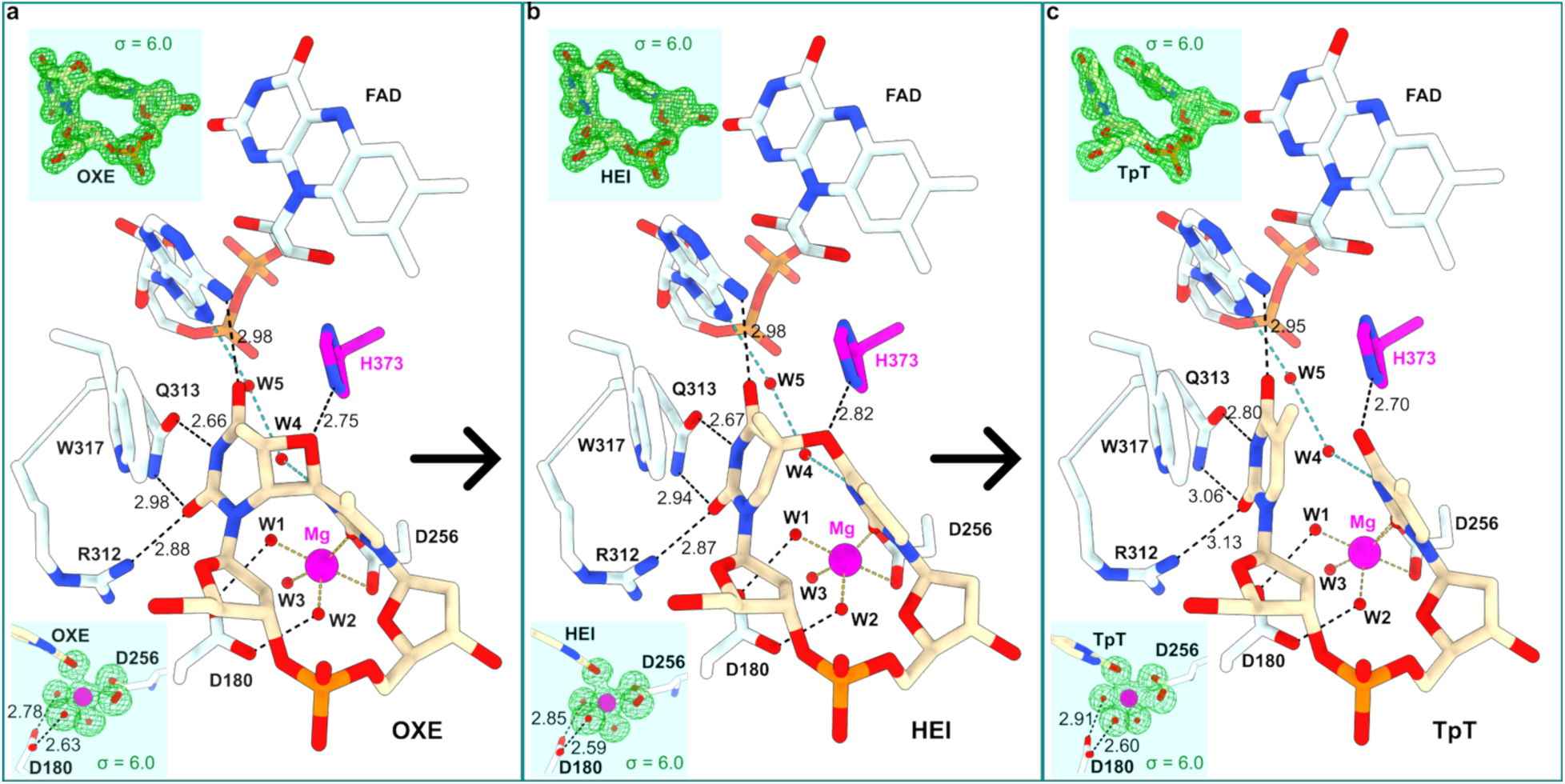
Sequential structures showing the interactions in the active sites containing: a, OXE; b, HEI; or c, TpT. The corresponding refined structures were resolved from the continuous-irradiation-experiment data (0.019, 0.075, and 0.717 MGy) and deposited under PDB codes 31TN (1.40 Å), 31UD (1.40 Å) and 31IF (1.51 Å), respectively. Dashed black and cyan lines represent hydrogen-bonding interactions. Gold dashed lines highlight the inner-sphere coordination (2.0–2.2 Å) of the hexa-coordinate Mg^2+^. The inset maps (green) are polder maps contoured at 6 σ, calculated with either the lesion model (OXE, HEI or TpT) or the Mg^2+^-centered-cluster model omitted.

### Identification of a hemi-cleaved intermediate along the X-ray driven repair pathway

Additionally, our experiments revealed a second intermediate (Fig. 3b) that cannot be described as either OXE or repaired TpT, or a mixture of both states. While the density of the 3′ ring appears planar and resembles that of an intact pyrimidine ring, the 5′-ring density is inconsistent with a planar aromatic ring, and therefore cannot be interpreted as a fully repaired thymine (Fig. S9-S10). When compared to OXE, the calculated polder map (Fig. 3b inset and S9) indicates that the oxetane C6-C4′ bond has been cleaved. However, the C5-O4′ bond still exists and the density around the C5 atom of the 5′ ring still forms a chiral center, a structural feature shared with 6-4PP and OXE. Therefore, we refer to this state as the hemi-cleaved intermediate (HEI), as it represents an asymmetric stepwise bond-breaking step of OXE that occurs before complete separation of the bases. In this structure (PDB: 31UD), the interbase C6-C4′ bond has opened, whereas the remaining interbase C5-O4′ linkage and the C5 chirality have not yet reached the product state. Despite these changes, all notable active-site interactions are preserved during the transition from OXE to HEI (Fig. 3).

### Intact thymine pair as the final state

With further increasing X-ray dose in both continuous-irradiation and pre-exposure experiments, we observed well-separated electron density signals on the 5′ and 3′ rings that match the geometry expected for a repaired thymine pair (Fig. 2, S9-S13). A calculated polder map (Fig. 3c) supports unambiguous assignment of two intact thymine nucleotides with planar pyrimidine rings, demonstrating that the X-ray-induced reaction results in genuine 6-4PP repair chemistry. In this final state, the repaired product (TpT) exhibits planar pyrimidine rings while still maintaining almost the same interactions with the active-site environment as those in the structures containing 6-4PP, OXE or HEI. The six-coordinate Mg^2+^ cluster is still highly ordered in this structure, where the DNA is repaired but still bound in the protein active site.

Interestingly, our high-resolution structures revealed an asymmetric contraction (0.1–0.2 Å) of the hexa-coordinate Mg^2+^ cluster and a slightly increased H373---O4′ distance in the intermediate states as compared to 6-4PP and TpT states (Fig. 4k-4l and S15). Further, a structural water (W4) approaches the 3′ ring during the repair (Fig. 4m). These events may serve as molecular reporters of the charge distributions on the lesion. Together, these observations suggest that efficient 6-4PP repair relies not only on a precisely orchestrated sequence of chemical transformations within the lesion itself, but also on coordinated reorganization of the surrounding electrostatic and hydrogen-bonding network, allowing the active site to adapt continuously while preserving its overall structural integrity.

**Fig. 4.**
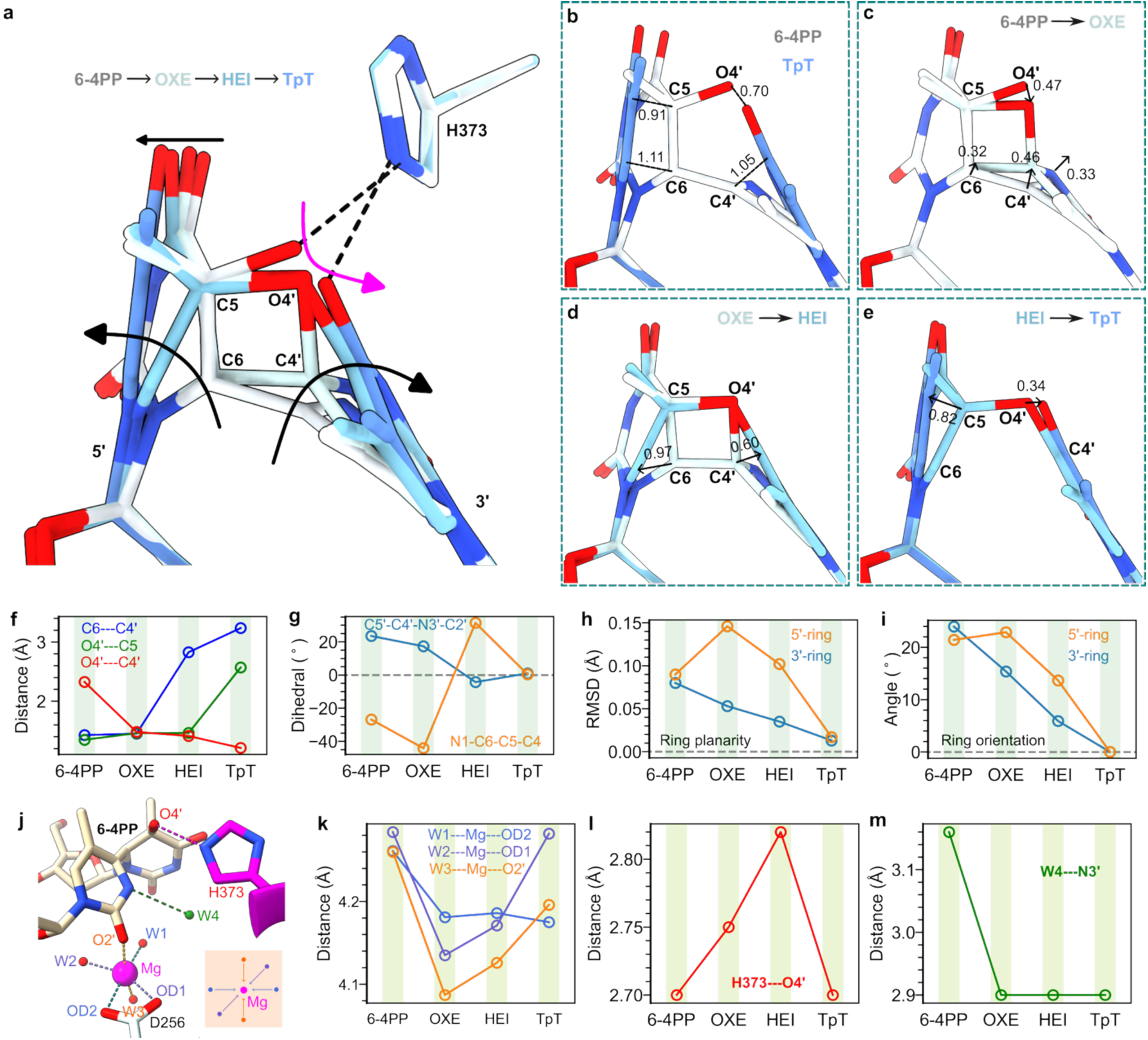
Active-site atomic displacements. **a,** Structural overlay of four lesion states reveals that during DNA repair, both the 5′ and 3′ rings rotate towards the product (TpT) state and the transferred O4′ moves along an arc-like trajectory pivoted around the Nε2 atom of H373. **b,** Geometrical differences between 6-4PP and TpT. **c-e,** Structural transitions between sequential reaction states. Distances shown in (b-e) are in Å. The overlays (a-e) were obtained by aligning the FADs in the structures (PDBs: 31TN, 31UD and 31IF) containing OXE, HEI and TpT to the FAD in the structure (PDB: 31JN) containing 6-4PP. **f,** Oxetane bond formation and breakages measured by the distances between oxetane-ring atoms. **g,** Dihedral angles showing pyrimidine ring distortion. **h,** Ring planarity measured by the RMSD distance from each reaction state to the corresponding best-fit planar ring. **i,** Relative orientations of the pyrimidine rings compared to the repaired state (TpT). **j,** Snapshot of active-site environment. **k,** Contraction of Mg^2+^-centered cluster during 6-4PP repair. The dimensions of the oxygen shell were quantified by the sum of distances from Mg^2+^ to two opposite oxygen atoms. **l,** Distance from the Nε2 atom of H373 to the transferred O4′ atom. **m,** Distance of a water molecule (W4) to the N3′ atom of the 3′ ring.

## Discussion

We report a sequence of four species in a (6-4) photolyase, ultimately terminating in intact, repaired DNA (Fig. 4). As our experiments employed X-rays to initiate this repair chemistry, as opposed to the blue light that is operative under natural conditions^9^, a key question is if our intermediates are also populated in blue-light-driven repair. During light-driven repair, photon absorption produces an excited FADH⁻, which in turn transfers an electron to the lesion^9^. By contrast, in our experiments, X-rays generate photoelectrons, secondary electrons and radicals in macromolecular crystals^50,51^. Such electrons can reduce flavins, including FAD in CPD photolyase crystals^36^, and moreover were observed to generate repaired DNA in the active site of a CPD photolyase^37^. Electrons liberated by X-rays may therefore reduce 6-4PP directly or indirectly via FAD, after which physiological reaction steps could proceed.

Three observations support, without proving, that the intermediates we report are generated by enzyme-coordinated redox chemistry rather than generic radiation damage. First, repair initiates after doses of just 2 kGy, and up to 0.7 MGy we observe no detectable global or site-specific damage: the protein, FAD, [4Fe-4S] cluster, and active-site hydrogen-bonding network, especially the H373—O4′ interaction, remain ordered (Fig. 3, S10, S12-S14). Second, in wild-type enzyme incorporating the antenna DLZ, accumulation of OXE and HEI is notably slower than in E38W (Fig. S16-S18). While more than 60% of lesions were fully repaired in E38W at 0.3 MGy, in wild type no apparent TpT was observed up to 0.8 MGy (Tables S4 and S5). We speculate DLZ may scrub free electrons or quench excited FAD; it similarly slows FAD photoreduction during blue-light irradiation (Fig. S3). Third, OXE provides direct structural evidence supporting long-standing models^10^ that invoke oxetane formation assisted by a conserved histidine, while OXE and HEI together provide a chemically plausible trajectory from 6-4PP to TpT (Fig. 4). We therefore hypothesize that X-rays replace physiological photoexcitation by supplying reducing equivalents, after which repair follows the enzyme’s intrinsic pathway. Our structures make the observed 6-4PP repair model precise and testable but do not definitively confirm its physiological relevance.

Our high-quality structures provide a step-by-step view of 6-4PP repair (Fig. 4a-4e). Overall, the pyrimidine rings reorganized their geometry significantly to minimize the displacement of the O4′ atom, probably designed to promote successful hydroxyl (O4′) reversal. The geometrical differences between 6-4PP and TpT (Fig. 4b) were reconciled by a stepwise mechanism: (1) (partial) transfer of O4′; (2) C6–C4′ breakage and restoration of a planar 3′ ring; and (3) C5–O4′ breakage and restoration of 5′ pyrimidine ring. The major, likely catalytically relevant, movement (∼0.47 Å) of O4′ occurred in OXE formation and was accompanied by the movement (∼0.46 Å) of C4′, reducing the O4′---C4′ distance from the initial 2.33 Å to a bonding distance of 1.48 Å (Fig. 4c). The first bond (C6–C4′) breakage was associated with significant shifts of C6 (∼1.0 Å) and C4′ (∼0.6 Å) in the opposite directions (Fig. 4d), which increased the C6---C4′ distance from 1.45 Å to 2.83 Å and might help stabilize the transiently formed radicals after C6–C4′ cleavage. Besides, the 3′ ring was already relaxed to a planar configuration and did not undergo significant conformational changes afterwards. In the final step, the splitting of C5–O4′ fully relaxed the chiral C5 atom, restoring the 5′ ring to a typical planer configuration (Fig. 4e).

Our results shed light on the most challenging event in 6-4PP repair: transfer of the hydroxyl group (-OH) between DNA bases. The H373—O4′ hydrogen bond persists through 6-4PP, OXE, HEI and TpT despite substantial chemical rearrangement (Fig. 1c and 3), making bidirectional proton transfer chemically accessible. O4′ follows an arc centered around H373 (Fig. 4a), suggesting that this conserved residue acts as both proton donor/acceptor and geometric pivot that constrains O4′ motion along the reaction coordinate, consistent with the concept that enzyme active sites steer reactive atoms along preferred trajectories^52,53^. This provides a separate explanation as to why the conserved histidine is indispensable^28^ for catalysis: it likely guides O4′ transfer throughout repair instead of letting O4′ dissociate and relocate freely. Besides, our structures indicate that the hydrogen atom (see Fig. 1a) of the transferred -OH could be transferred directly from O4′ to the N3′ atom of the 3′ ring without the involvement of intervening molecules^20^, because O4′ is only 2.44 Å from N3′ in OXE. What’s more, while the 3′ ring transformed progressively towards a pyrimidine, the 5′ ring seemed more distorted in OXE (Fig. 4g-4i). We suspect that the low quantum yields (∼0.1) of (6-4) photolyases^12^ reflect constraints imposed by the challenging repair chemistry itself rather than insufficient evolutionary optimization, and these enzymes adopt a delicate histidine-guided repair strategy to maximize the efficiency of converting photon energy into DNA repair.

In addition, our findings call for a renewed investigation into the mechanism of DNA repair by (6-4) photolyases. The HEI structure provides the first experimental evidence for the order of oxetane opening, which most models do not specify^10^. Contrary to the intuitive expectation that C5–O4′ breaks first, the HEI model suggests that C6–C4′ cleavage precedes C5–O4′ rupture. This order of bond breakage is consistent with the oxetane geometry (Fig. 4d): any species with an intact C6–C4′ bond but broken C5–O4′ is perhaps sterically strained. None of our datasets showed the converse species, although a very short-lived state cannot be excluded. Future QM/MM studies should therefore investigate the role of HEI, allowing mechanistic calculations to be constrained by the observed structural pathway rather than by hypothetical bond-cleavage sequences.

We further asked if our structures could inform us as to how Vc64PL specifically recognizes and binds to 6-4PP lesions. The protein:DNA interactions found in the 6-4PP lesion state are largely preserved in the repaired TpT (Fig. 1c and 3c). Therefore, the structural complementarity between the DNA lesion and the active site alone is unlikely to account for preferential lesion binding. We speculate that specificity may be established during the route into the active site, while the final bound pocket appears capable of accommodating both the substrate and product. Such an arrangement could facilitate efficient catalysis while avoiding excessively tight product binding, thereby promoting product release after repair.

Crucial questions remain regarding the mechanism of DNA repair by (6-4) photolyases, including the details of the photochemistry and the number of reducing equivalents required, the precise identity of all intermediates populated in light-driven repair and the role of iron-sulfur clusters in bacterial enzymes. These questions have been intractable over more than thirty years of study. We identified two intermediates bound to Vc64PL and determined the high-resolution structures of each. These structures provide a new foundation on which to propose and incisively test models of the repair chemistry, perhaps finally fulfilling the long-standing quest for the (6-4) photolyase mechanism.

## Methods

### Protein and DNA substrate preparation

Protein was produced by slightly modifying published conditions^40,43^. The amino acid sequences of Vc64PL (UniProt Q9KLD7) and Vc64PL-E38W were sent to BioCat (Heidelberg, Germany) for codon optimization and gene synthesis. The codon-optimized genes were cloned into pET-28a between the NdeI and BamHI restriction sites to generate an N-terminal His6-tagged construct. Briefly, recombinant proteins were expressed in *Escherichia coli* BL21(DE3) cells (Invitrogen) using auto-induction Terrific Broth medium (Formedium, UK) following reported expression conditions^43^. Then the proteins were purified by sequential Ni-affinity, heparin-affinity and size-exclusion chromatography (Cytiva) following a reported purification protocol^40^. Purified protein was concentrated to approximately 20 mg/mL, exchanged into storage buffer (20 mM Tris-HCl, 200 mM NaCl, 20% (v/v) glycerol, pH 7.6), flash-frozen in liquid nitrogen and stored at −80 °C until use.

The (6-4) photoproduct (6-4PP) substrate was synthesized by ultraviolet irradiation of the oligonucleotide d(GTATTATG) following a published protocol^35^. This resulted in an 8-mer substrate containing a T<6-4>T lesion in the center of the DNA strand.

### Crystallization

Protein-DNA complexes were prepared by mixing purified protein with lesion-containing DNA in a molar ratio of ∼1:1.5 immediately before crystallization. Initial crystallization screening was performed at the EMBL Hamburg Sample Preparation and Characterisation (SPC) Facility. Optimized sitting-drop vapor-diffusion conditions yielded two crystal forms. Disk-shaped crystals (space group P6422) grew from 0.1 M HEPES, 2 M ammonium sulfate, 4% (v/v) PEG 400, 1 mM MgCl2 and 10% (v/v) glycerol (pH 7.5). Rod-shaped crystals (space group P212121) were obtained from 0.1 M HEPES, 20% (w/v) PEG 5000, 10 mM MgCl2 and 10% (v/v) glycerol (pH 7.5). For each condition, 1 µL protein-DNA complex (20 mg/mL for disk-shaped crystals, 15 mg/mL for rod-shaped crystals) was mixed with 1 µL crystallization solution. Crystals typically appeared in 1–2 days at room temperature. All single crystals were flash-cooled directly in liquid nitrogen without additional cryoprotection.

Microcrystals for serial crystallography were produced by seeded batch crystallization. Initial seed stock was prepared by crushing single disk-shaped crystals using 2-mm plastic beads. 300 µL of protein-DNA complex (10 mg/mL) was mixed with 600 µL of crystallization solution containing 0.1 M HEPES, 2 M ammonium sulfate, 4% (v/v) PEG 400, and 1 mM MgCl2 (pH 7.5), supplemented with 1 µL seed solution. The suspension was incubated with constant shaking (400 rpm) at 16 °C (Eppendorf ThermoMixer C). Microcrystals reached a typical size of approximately 10 µm after 12 hours.

### Serial femtosecond crystallography

Serial femtosecond crystallography data were collected from dark-state E38W:6-4PP microcrystals at the Alvra endstation of the SwissFEL Aramis beamline at the Paul Scherrer Institute, Switzerland. Diffraction data were collected at room temperature using 11.98 keV X-ray pulses (40 fs duration, 100 Hz repetition rate, approximately 990 µJ energy) and a JUNGFRAU 16M detector. The incident pink beam was attenuated to 30% transmission and focused to a spot size of ∼3.5 × 2.5 µm^2^ (FWHM). Using two glass syringes and a coupling device, a slurry of microcrystals was mixed in volumetric ratio of 1:3 with 22% w/v hydroxyethyl cellulose (Sigma-Aldrich) swelled in water and delivered to the X-ray interaction region using a high-viscosity injector^54^ via a needle with an inner diameter of 75 µm.

Diffraction images were processed with CrystFEL (v0.12.0)^55^. Bragg peaks were identified using Peakfinder8 (as implemented in Cheetah^56^) with a signal-to-noise ratio threshold of 5 and an intensity threshold of 1200 ADU. Diffraction patterns were indexed with XGANDALF^57^. Peak integration was performed using the rings-grad method with integration radii of 3, 5 and 7 pixels for the inner, middle and outer radii, respectively. Integrated intensities were merged and scaled with *partialator*^55^ using the unity model with five refinement cycles. The final dataset was merged in space group P6422 to a resolution of 1.67 Å using a CC1/2 value of 0.5. Data collection and refinement statistics are summarized in Tables S1.

### Dose-resolved crystallography

An initial dataset was collected from a rod-shaped crystal (approximately 400 × 80 × 50 μm^3^, space group P212121) at beamline P14 of PETRA III (DESY, Hamburg, Germany) using 12.8 keV X-rays and an EigerX 16M detector at 100 K. This crystal was illuminated with a 300 µm top-hat beam at 100% transmission (8.8 × 10^12^ photons/second or 1.2 × 10^8^ photons/second/µm^2^). The data were collected with 0.1° oscillation per image (integration time 7.5 milliseconds) and yielded a structure (PDB: 31IF) containing a fully repaired DNA shown as TpT in Fig. 3c. A follow-up continuous-irradiation experiment was performed at P14 using the same beamline setup. To better disentangle mixed reaction states, disk-shaped crystals (approximately 300 × 300 × 80 μm^3^, space group P6422) were used and the X-ray transmission was gradually increased from 1% and 40%. Each crystal was only used once to collect one dataset (one rotation or 3600 images) at a selected X-ray transmission to avoid radiation contamination. 360°-rotation datasets were collected with 0.1° oscillation per image (integration time 7.5 millisecond) and subsequently divided into 80° wedges with 40° overlap between adjacent wedges for dose-dependent structural analysis.

Pre-exposure experiments were performed at the MicroMAX beamline of MAX IV (Lund, Sweden) using a 200 µm top-hat X-ray beam (13.7 keV, 1.3 × 10^14^ photons/second or 4.1 × 10^9^ photons/second/µm^2^, ΔE/E ∼ 1%) and a JUNGFRAU 9M detector at 100 K. Rod-shaped crystals (approximately 200 × 80 × 50 μm^3^, space group P212121) were subjected to various pre-exposure times (no pre-exposure, 50-milliseconds pre-exposure and 1-second pre-exposure) followed by a 60-second dark waiting period before data collection was performed (180° rotation with 0.1° oscillation per frame, integration time 0.5 millisecond). The crystals were subjected to continuous rotation throughout pre-exposure, waiting and data collection. The beamline shutter (Cedrat) was programmed and controlled by PandABox to produce a timing sequence for pre-exposure and waiting intervals.

For all crystals, the longest axis was aligned to be parallel to the rotation axis of the beamline goniometer. All crystals used in these experiments were prepared in an oxidized state with no reducing agent added. All rotational datasets were processed and scaled with DIALS (v3.25.0)^58^. Resolution limits were selected using a CC1/2 value of 0.5. Data collection and refinement statistics are summarized in Tables S2 and S3.

Absorbed X-ray doses were estimated with RADDOSE-3D (v5.0)^48^ using the measured crystal dimensions, beam profile, photon energy, flux, exposure time, rotation range and macromolecular composition. Average Diffraction Weighted (ADW) dose was used throughout to quantify radiation exposure. Two RADDOSE-3D input files are provided at the end of the Supplementary Information to facilitate reproduction of the dose calculations.

### Structure solution and refinement

The Vc64PL-E38W:6-4PP structure (PDB: 31JN) was determined by molecular replacement with Dimple (from the CCP4 suite^59^) using the PDB entry 8A1H^40^ as the search model. All subsequent complex structures were solved by molecular replacement using 31JN as the initial model. Model building was performed using Coot^60^ and refinement was performed using Phenix^49^. Protein structures and electron-density maps were visualized via UCSF ChimeraX^61^.

#### 6-4PP

Initial coordinates (64T, 5PY) were extracted from the crystal structure of *Drosophila melanogaster* (6-4) photolyase (PDB entry 3CVU)^17^, converted to MOL2 format using Avogadro^62^, and submitted to the Grade Web Server (https://grade.globalphasing.org) to generate initial geometry restraints. Initial refinement revealed that several automatically generated restraints were incompatible with the experimental electron density. In particular, planarity restraints involving the deoxyribose rings were removed, and restraints describing C=N and C=O double bonds, pyrimidine-ring planarity and torsional distortions around the chemically perturbed C6 atom of the 5′ pyrimidine and C4′ atom of the 3′ pyrimidine were relaxed to allow the ligand geometry to be determined predominantly by the SFX data. The resulting restraint dictionary provided excellent agreement between the refined model and the experimental electron density and was used for refinement of all structures containing the 6-4PP lesion. To maintain consistency with previously deposited structures^17,18^, our model (PDB: 31JN) represent the 6-4PP as two linked residues: 64T and 5PY.

#### OXE and HEI

All data wedges from the continuous-irradiation experiment were initially refined against our SFX structure (PDB: 31JN). Subsequently, polder maps were calculated for all data wedges with the lesion model omitted; and data wedges that appeared to be dominated by OXE or HEI were selected for refining the intermediate models. Preliminary coordinates for OXE and HEI were generated by manually modifying the refined 6-4PP model. Initial coordinates and restraint dictionaries for OXE and HEI were generated using the Grade Web Server (https://grade.globalphasing.org) and used for preliminary refinement of the selected data wedges of the corresponding intermediates. The resulting refined geometries were subsequently used to generate the final restraint dictionaries, which were again used to refine the selected data wedges to ensure that all restraints of OXE or HEI were appropriate. We performed intermediate (OXE, HEI) refinement and occupancy refinement (see below) iteratively to select the data wedges (at 0.019 and 0.075 MGy) that have the highest occupancies of OXE or HEI. These data wedges were used in a final round of refinement to obtain the deposited models (PDBs: 31TN, 31UD) containing OXE and HEI. OXE and HEI were deposited as individual ligands and assigned the ligand identifiers: A1KCH and A1KBV, respectively, by the Protein Data Bank. The obtained restraints of OXE and HEI were also used in refining data from the pre-exposure experiment.

Standard Phenix^49^ restraint dictionaries were used for all remaining nucleotides, cofactors and protein residues, including DA, DT, DG, FAD, DLZ and the [4Fe–4S] cluster. TpT was modelled as two adjacent DT monomers.

Dose-resolved datasets frequently contained mixtures of multiple lesion states. These were modelled as alternative conformations (6-4PP/OXE, OXE/HEI, or HEI/TpT) with occupancies constrained to sum to unity. During occupancy refinement, atomic coordinates were fixed and isotropic B factors of the atoms of the modelled reaction states were set to 20 Å^2^ so that only their occupancies were refined. Occupancies were refined using Phenix^49^ and monitored for 10 refinement cycles to ensure convergence. Note that separate restraint dictionaries were made for both 6-4PP and TpT so that each state could be modelled/refined as an alternative conformer of OXE or HEI. The resulting occupancies are summarized in Table S5.

### Structural geometry analyses

Structural changes (Fig. 4a-4i) accompanying 6-4PP repair were quantified from the refined coordinates of the 6-4PP (PDB: 31JN), OXE (PDB: 31TN), HEI (PDB: 31UD) and repaired TpT (PDB: 31IF) structures. Prior to geometric analysis, each structure was superposed onto 31JN by least-squares fitting of the C/N/O atoms of FAD. The RMSD values of FAD alignments are 0.0075, 0.0077 and 0.0085 Å, respectively, for OXE, HEI and TpT.

Atomic displacements between adjacent reaction states (6-4PP→OXE, OXE→HEI and HEI→TpT) were calculated from the refined atomic coordinates following FAD-based structural superposition.

Local distortion of the 5′ pyrimidine ring was quantified using the assigned dihedral angle N1–C6–C5–C4, which reports both the magnitude and direction of out-of-plane distortion around the chemically perturbed C6 atom. Likewise, local distortion of the 3′ pyrimidine ring was quantified using the assigned dihedral angle C5–C4′–N3′–C2′, describing the local geometry around the C4′–N3′ bond. Values approaching 0° indicate restoration of a locally planar pyrimidine geometry.

The planarity of each pyrimidine ring was evaluated by fitting a least-squares plane through the six endocyclic ring atoms (N1, C2, N3, C4, C5 and C6). Ring distortion was quantified as the root-mean-square deviation (RMSD) of these atoms from the corresponding best-fit plane.

To quantify the overall reorientation of the pyrimidine rings during repair, best-fit planes were determined for each pyrimidine ring. The angle between the plane normal of each intermediate and that of the corresponding repaired thymine ring was then calculated to quantify progressive restoration of the repaired DNA geometry.

## Data availability

Data supporting the findings of this study are available within the Article, the Supplementary Information or the source data. Structural models, along with the structure factors presented here, can be found under PDB accession codes: 31NN, 31JN, 31IF, 31TN, 31UD, 31TQ, 32AM, 32AN, 32AO, 32SG, 31JD, 32UC, 32UD.

## Acknowledgements

This work was supported by the Cluster of Excellence “CUI: Advanced Imaging of Matter” of the Deutsche Forschungsgemeinschaft (DFG) – EXC 2056 – project ID 390715994. Additional support was provided by the Helmholtz Association through a Young Investigator Group (YIG) award to T.J.L. and by the National Natural Science Foundation of China (Grants T2495220 and T2495222 to D.Z.).

XFEL experiments were performed at the Alvra endstation of the Aramis beamline at SwissFEL, Paul Scherrer Institute (Proposal 20250954). We thank Dr. Camila Bacellar and Dr. Claudio Cirelli for their help in beamline steup. Continuous-radiation crystallography experiments were carried out at the P14 beamline of PETRA III. We thank Dr. Eike Schulz for providing beamtime (Proposal MX-1021). Pre-exposure experiments were performed at the MicroMAX beamline of MAX IV (Proposals 20261161 and 20261292). MicroMAX is funded by the Novo Nordisk Foundation (Grant NNF17CC0030666). We thank Dr. Mirko Milas and Dr. Sofia Kapetanaki for their help in beamline setup. We acknowledge the MAX IV Laboratory for beamtime on the MicroMAX beamline under proposal 20261161 and 20261292. Research conducted at MAX IV, a Swedish national user facility, is supported by Vetenskapsrådet (Swedish Research Council, VR) under contract 2018-07152, Vinnova (Swedish Governmental Agency for Innovation Systems) under contract 2018-04969 and Formas under contract 2019-02496. MicroMAX is funded by the Novo Nordisk Foundation under the grant number NNF17CC0030666.

We thank Dr. David Ruiz Carrillo for his assistance in crystallization screening in the Sample Preparation and Characterisation (SPC) Facility (EMBL Hamburg) and Dr. Robin Schubert for his assistance in sample preparation in the EuXFEL Biology and Chemistry User Labs. The authors thank Dr. Sebastian Günther and Dr. Patrick Reinke (P09, PETRA III), Prof. Arwen Pearson and Dr. David von Stetten (P14.2 T-REXX, PETRA III, access via the mx1008, TREXX4 and TREXX7 BAGS), Dr. Shibom Basu, Dr. Samuel Rose, and Dr. Daniele de Sanctis (ID29, ESRF, access via bag proposal MX-2727), Dr. Dean Lang, Dr. Manoop Chenchiliyan, Dr. Monik Bjelcic, and Dr. Cecilia Casadei (MicroMAX, MAX IV, access via proposal 20252324) for their help in testing crystal diffraction quality.

C.Y. thanks Prof. Dusan Turk, Dr. Sebastian Günther, Dr. Patrick Reinke, Dr. Chufeng Li and Mr. Bjarne Klopprogge for insightful discussions. C.Y. gratefully acknowledges Dr. Dominik Oberthuer for invaluable suggestions and training in macromolecular crystallography.

## Author contributions

C.Y., H.N.C., D.Z. and T.J.L. contributed to project conception. C.Y. designed and led the structural study, performed the experiments, analyzed and interpreted the data, and prepared the figures. Q.Y. and X.C. prepared the DNA substrates and performed spectroscopic experiments. C.Y. and J.N. conducted MAX IV beamtime. C.Y., E.V.B. and T.J.L. conducted SwissFEL beamtime. C.Y. and T.J.L. wrote the manuscript. S.F., J.N., E.V.B., H.N.C. and D.Z. edited the manuscript. H.N.C., D.Z. and T.J.L. acquired funding and provided resources. T.J.L. supervised the research. All authors discussed the results and approved the final manuscript.

