## Supplementary Information for "The structures of two intermediates *en route* to DNA repair within a (6-4) photolyase"

### for

This file contains:

Supplementary Discussions 1-2

Supplementary figures Fig. S1-S19

Supplementary tables Table S1-S6

Supplementary RADDOSE-3D input files

Supplementary references

### Supplementary Discussions

#### 1. Functional validation of Vc64PL-E38W

*Photoactivity of Vc64PL-E38W*

To verify that removal of the antenna chromophore did not compromise the catalytic function of Vc64PL, we examined the photochemical activity of the E38W mutant in solution.

Absorption spectra (Fig. S3) of purified wild-type (WT) and E38W proteins exhibited the characteristic spectral features of the catalytic FAD cofactor and the [4Fe–4S] cluster, consistent with previous spectroscopic studies of prokaryotic (6-4) photolyases[1,2]. As expected, the WT protein displayed stronger absorption in the near-UV region because of the presence of the antenna chromophore DLZ.

Photoreduction experiments were performed using WT and E38W at a concentration of 75 μM in the presence of 20 mM DTT under continuous blue-light illumination (470 nm, LIU470A from ThorLab). Both proteins underwent progressive photoreduction, as indicated by the characteristic decrease in the absorption bands of oxidized FAD. Difference spectra (Fig. S3) closely resembled those reported for flavin photoreduction in photolyases[3]. Under identical illumination conditions, photoreduction of E38W proceeded more rapidly than that of the WT protein.

The DNA repair activity of E38W was subsequently examined using purified oligonucleotides that contain a central 6-4PP lesion. Protein and DNA were mixed at a molar ratio of approximately 1:200 in the presence of 20 mM DTT and 10 mM $MgCl_2$. Upon blue-light illumination, the characteristic absorption band of the 6-4PP lesion at approximately 325 nm decreased progressively, accompanied by a corresponding increase in DNA absorption near 266 nm, demonstrating efficient repair of the UV lesion (Fig. S4). Because DNA repair by WT has been reported previously[4], only the E38W mutant was examined in these measurements.

Together, these results demonstrate that removal of the antenna chromophore does not abolish either flavin photochemistry or catalytic DNA repair, validating E38W as an appropriate model system for the structural studies presented in this work.

*DNA repair in crystals*

To determine whether E38W retained catalytic activity in the crystalline state, disk-shaped protein:DNA co-crystals were soaked overnight with approximately 20 mM DTT in the dark to allow diffusion of the reductant into the crystal lattice while minimizing premature photoactivation. The crystals were then either illuminated with blue light (470 nm, LIU470A from ThorLab) at 4 °C for 3 h, or retained in the dark as a control, before being flash-cooled in liquid nitrogen.

Diffraction analysis showed that the electron density corresponding to the bound DNA disappeared following blue-light illumination, whereas strong DNA density was retained in control crystals that were not illuminated (Fig. S5). These observations demonstrate that the protein remains catalytically competent in the crystalline state and that the crystal lattice does not prevent photoinduced repair of the DNA lesion.

The preservation of catalytic activity in both solution and crystals provides an essential foundation for interpreting the X-ray-induced structural changes described in the main text as bona fide intermediates of the enzymatic repair reaction.

### 2. Wild-type Vc64PL and E38W show different repair kinetics

High-energy X-rays generate photoelectrons, secondary electrons and reactive radical species throughout protein crystals[5,6]. Therefore, in principle, the observed DNA repair could arise either from nonspecific radiation chemistry or from activation of the native enzymatic repair mechanism.

The presence of the antenna chromophore DLZ provides a useful mechanistic probe because it is expected to influence flavin photochemistry while contributing negligibly to X-ray absorption owing to the absence of heavy atoms. Consequently, both proteins should generate similar populations of radiation-induced electrons under identical experimental conditions.

We conducted crystallographic experiment using a 300-μm WT disk-shaped crystal that grew from the same crystallization conditions as E38W. The diffraction data were collected under the same condition that led to the fully repaired DNA in E38W (PDB: 31IF), namely, full flux ($8.8 \times 10^{12}$ photons/second) of the P14 X-ray beam. Electron density corresponding to the OXE and HEI intermediates was observed sequentially in the WT crystal as the total exposure time increased, and the refined structures (PDBs: 32SG, 32UC, and 32UD) revealed essentially identical active-site interactions (Fig. S16) as those in E38W. These observations indicate that the same sequence of structural intermediates is formed irrespective of the presence of DLZ.

Despite the similar structural sequency, the overall repair efficiency differed markedly between the two proteins. Whereas E38W was dominated by the TpT state after 0.3 MGy X-ray radiation, WT crystals accumulated little repaired product even after 0.7 MGy X-ray radiation. Instead, increasing X-ray dose resulted in progressive degradation of the electron density for the 5′-thymine methyl group together with increasing disorder around the $Mg^{2+}$-coordinated octahedral cluster (Fig. S16-18). Consistent with these observations, diffraction quality deteriorated progressively from the substrate to the intermediate states in the WT datasets (Table S4), indicating that radiation damage accumulated more rapidly than DNA-repair intermediates or products.

Moreover, weak but reproducible mFo−DFc density was consistently observed around the DLZ chromophore in the WT data (Fig. S19). Although the signal was not sufficiently strong to justify modelling an alternative DLZ conformation, it is consistent with a subtle structural response of the antenna chromophore to X-ray irradiation and involvement of DLZ in X-ray-driven redox chemistry.

The major difference between WT and E38W is the presence of the antenna chromophore DLZ. Previous spectroscopic studies have shown that excitation energy is transferred efficiently between the antenna chromophore and the flavin cofactor in photolyases[7]. Because the WT crystals used in this study contained oxidized FAD, one possible explanation is that electronic excitation associated with X-ray-induced flavin activation is partially quenched through energy transfer to DLZ, thereby reducing the fraction of catalytically productive flavin. Consistent with this interpretation, E38W undergoes substantially faster flavin photoreduction than WT under blue-light illumination (Fig. S3). Alternatively, DLZ might act as a generic electron sink, accepting electrons that would otherwise directly reduce the 6-4PP lesion. More complex redox schemes, involving combinations of the iron-sulfur cluster, FAD, DNA bases, redox-active protein sidechains and solvent cannot be excluded.

One possible explanation for the X-ray-induced 6-4PP repair is that the FAD is excited by X-ray and it performs an oxidative 6-4PP repair reaction. The excited FAD abstracts an electron from 6-4PP, generating a positively charged lesion species and following a cation radical pathway that was investigated in an early QM study[8]. The bond splitting order observed in our work is consistent with a cation radical pathway[8]. However, so far, no oxidative DNA repair has been achieved in any photolyases under blue-light illumination. Considering that X-ray-induced FAD reduction[9] and DNA repair[10] have been observed before, a more plausible explanation is that in our crystals FAD was first reduced by X-ray and then the catalytic reduced flavin was activated by X-ray. We did not observe an apparently bended flavin cofactor, a characteristic feature of reduced FAD observed in other photolyases[9,11]. But a near planar reduced FAD might be a unique feature in prokaryotic (6-4) photolyases, as shown in a recent study in which the flavin bending due to reduction was tiny in a (6-4) photolyase from *Caulobacter crescentus*[2]. Further research is necessary to either prove or disprove the existence of an X-ray-driven oxidative repair mechanism.

Although the precise mechanism by which X-rays produce productive 6-4PP repair chemistry cannot be established, the different repair efficiencies observed for WT and E38W strongly suggest that the reaction is not driven solely by nonspecific radiation chemistry.

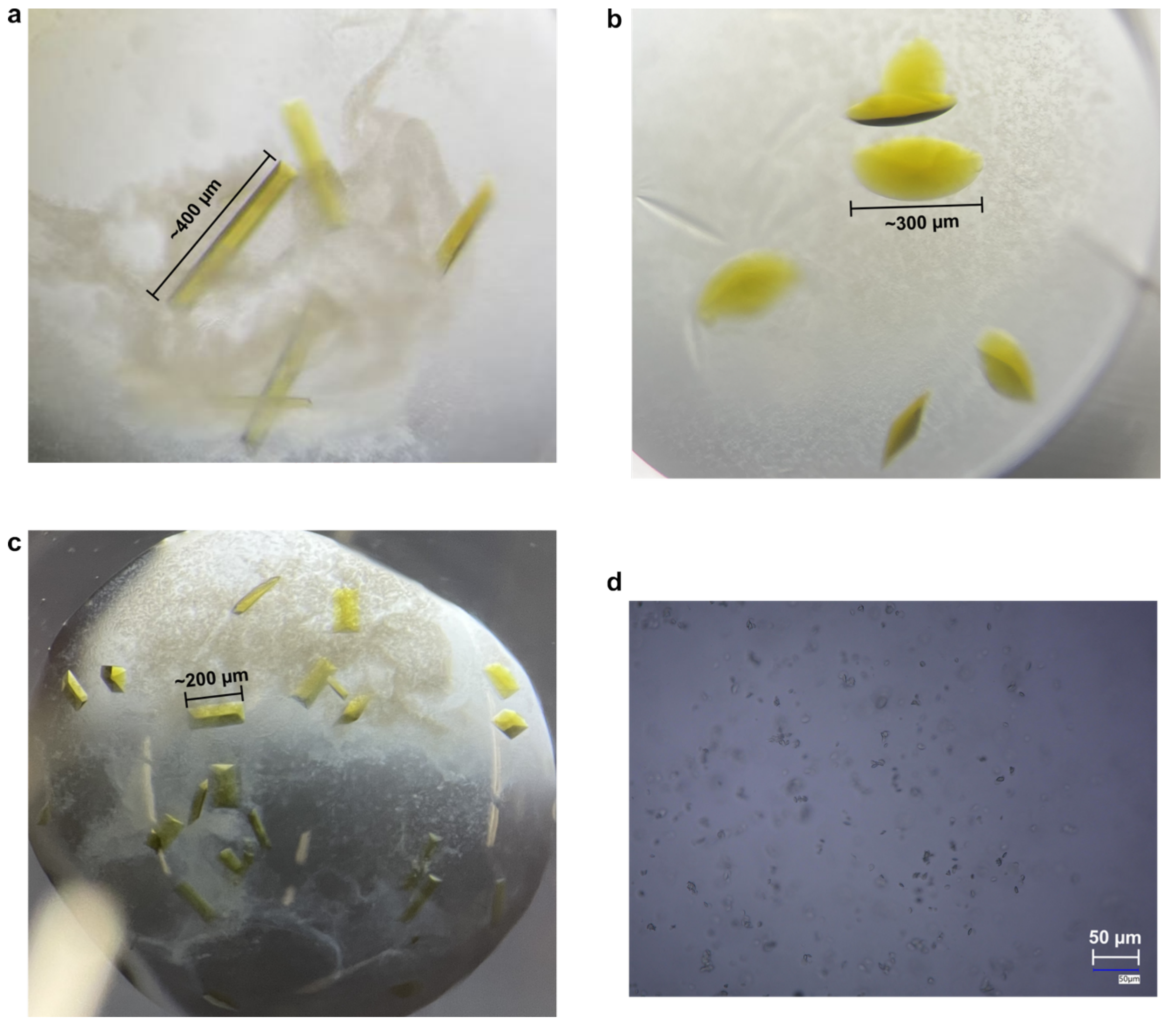


**Fig. S1 | Vc64PL-E38W+6-4PP co-crystals. a,** rod-shaped crystals (~400 × 80 × 50 μm$^3$) developed in early phase of this work. **b,** disk-shaped crystals (~300 × 300 × 80 μm$^3$). **c,** rod-shaped crystals (~200 × 80 × 50 μm$^3$). **d,** disk-shaped microcrystals (largest dimension ~10 μm). Note than rod-shaped crystals shown in **a** and **c** grew from the same crystallization solution by using different seeds.

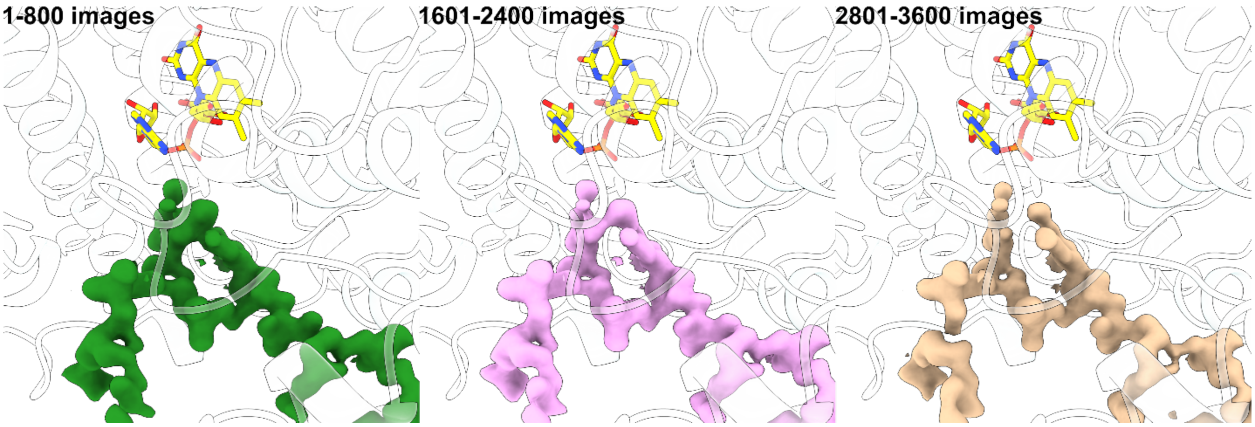


**Fig. S2 | Preliminary electron-density maps reveal 6-4PP repair intermediates.** Shown are the mFo-DFc maps (3 σ) from initial refinement against a DNA-free E38W model in which we first observed interesting X-ray-driven reaction intermediates. The data are from one rotational data collection from the same crystal (360° with an oscillation of 0.1°, integration time was 7.5 milliseconds, X-ray flux was $1.7 \times 10^{12}$ photons/second). Although these maps do not represent the final refinement results that we ultimately presented, they show obvious intermediate features from 6-4PP repair that inspired the present study. The flanking bases are also visually identifiable. For clarity, only density in the area expected to be occupied by the DNA was shown. The density was cropped using UCSF ChimeraX[12] by aligning the density map to a E38W:6-4PP complex model (PDB: 31JN).

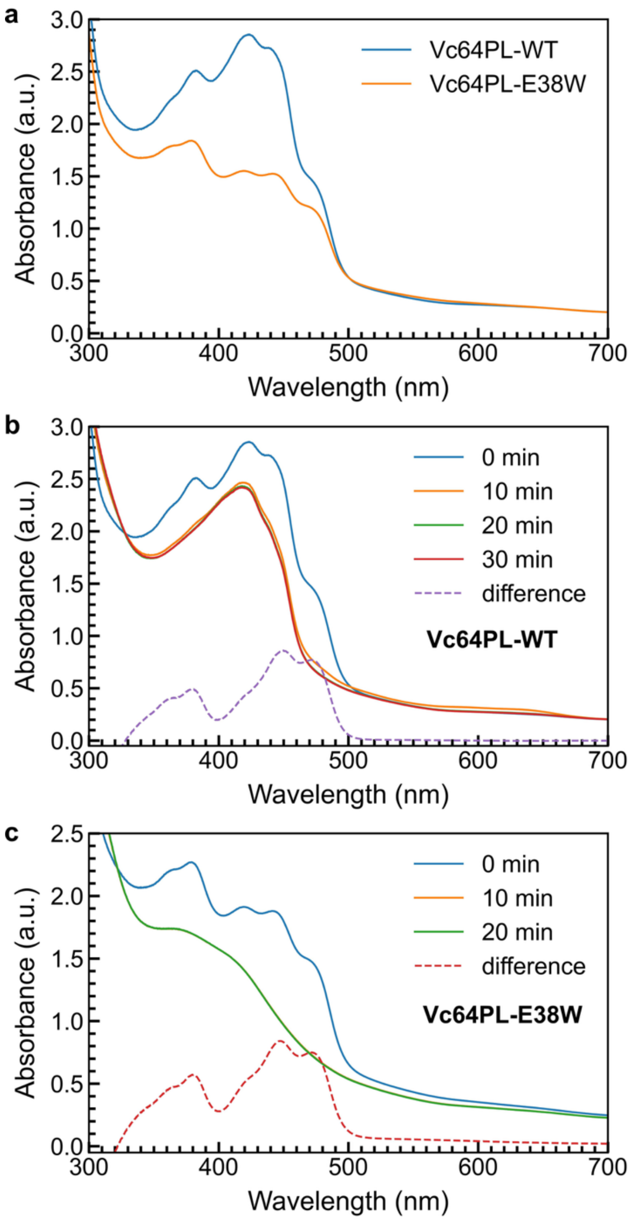


**Fig. S3 | Steady-state photoreduction. a,** UV-Vis absorption spectra of Vc64PL WT and E38W. The spectral difference is due to a lack of the DLZ cofactor in E38W. The spectra were re-scaled to align the absorbance from 500 to 700 nm for better visualization. **b** and **c** show the photoreduction of WT and E38W, respectively; both yielded difference spectra resembling the absorption of oxidized FAD bound in a protein environment. The protein concentration was estimated by calculating FAD concentration from the difference spectra. The same concentration (75 μM) was used for WT and E38W. Photoreduction of E38W is clearly faster than that of WT under the same conditions.

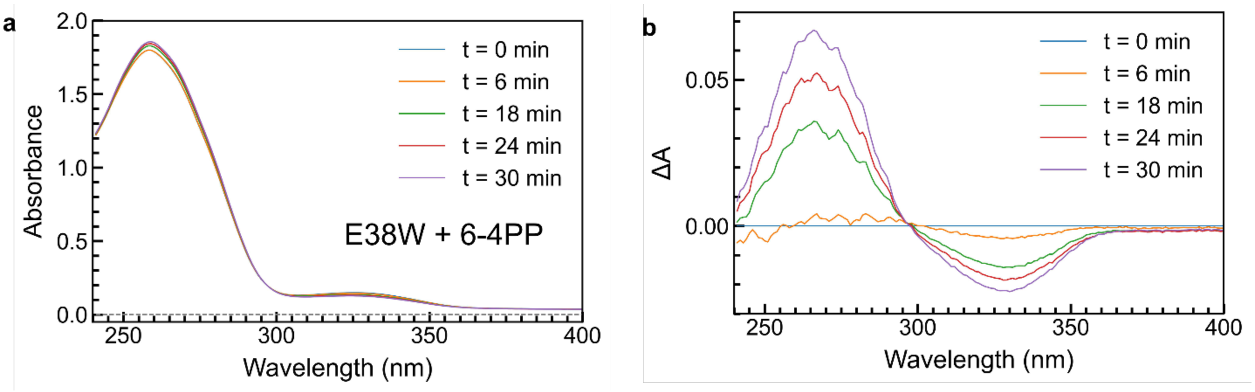


**Fig. S4 | Steady-state 6-4PP repair in solution.** The DNA-repair ability of E38W was confirmed by a steady-state repair experiment. **a,** absorption spectra measured during 6-4PP repair. **b,** difference absorption spectra obtained by subtracting the 0-min spectrum as background. 1 μM E38W was mixed with 200 μM 6-4PP-containing 8-mer DNA. In the presence of 20 mM DTT, 6-4PP was continuously repaired under blue-light illumination. The spectra show a decrease in absorption at 325 nm and an increase at 266 nm, which are characteristic of 6-4PP repair and thymine formation, respectively.

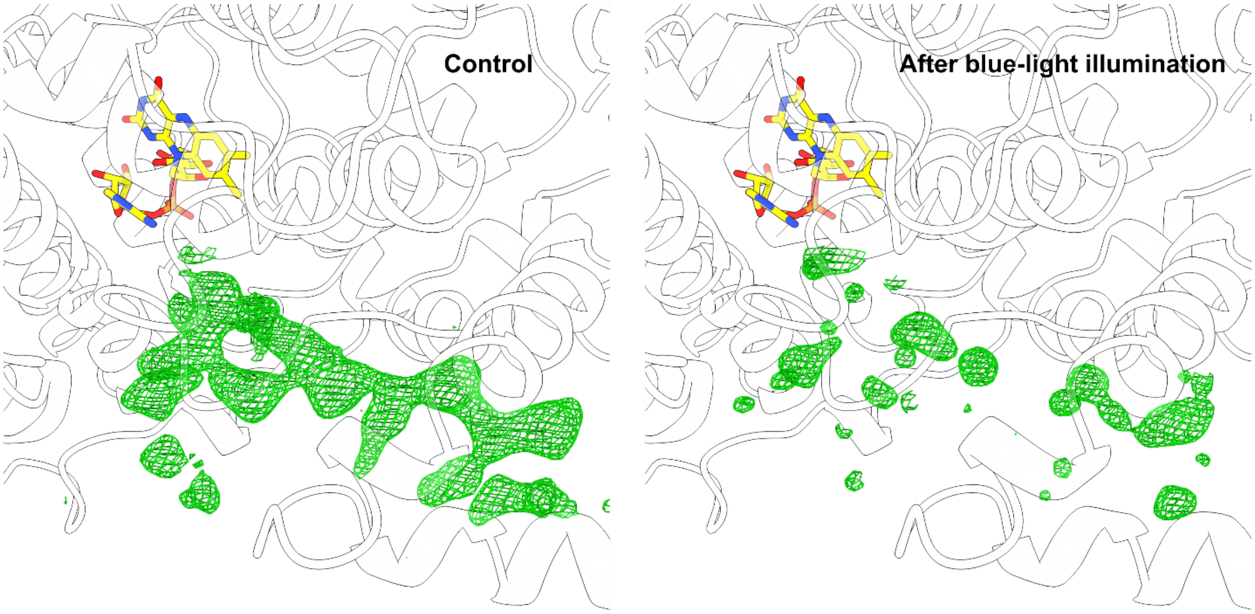


**Fig. S5 | Steady-state 6-4PP repair in crystal.** E38W:6-4PP complex (disk-shaped, ~100 μm in diameter) crystals were soaked with 20 mM DTT and then exposed to blue light at 4ºC for 3 hours. Presented are mFo-DFc maps (3 σ) obtained from refinement against a DNA-free E38W model. For clarity, only density in the area expected to be occupied by the DNA was shown. The density was cropped using UCSF ChimeraX[12] by aligning the density map to a E38W:6-4PP complex model. The decrease in observed density is consistent with 6-4PP repair and product release following blue light illumination, suggesting that our crystals are active and support the enzyme and DNA dynamics necessary to effect 6-4PP lesion repair.

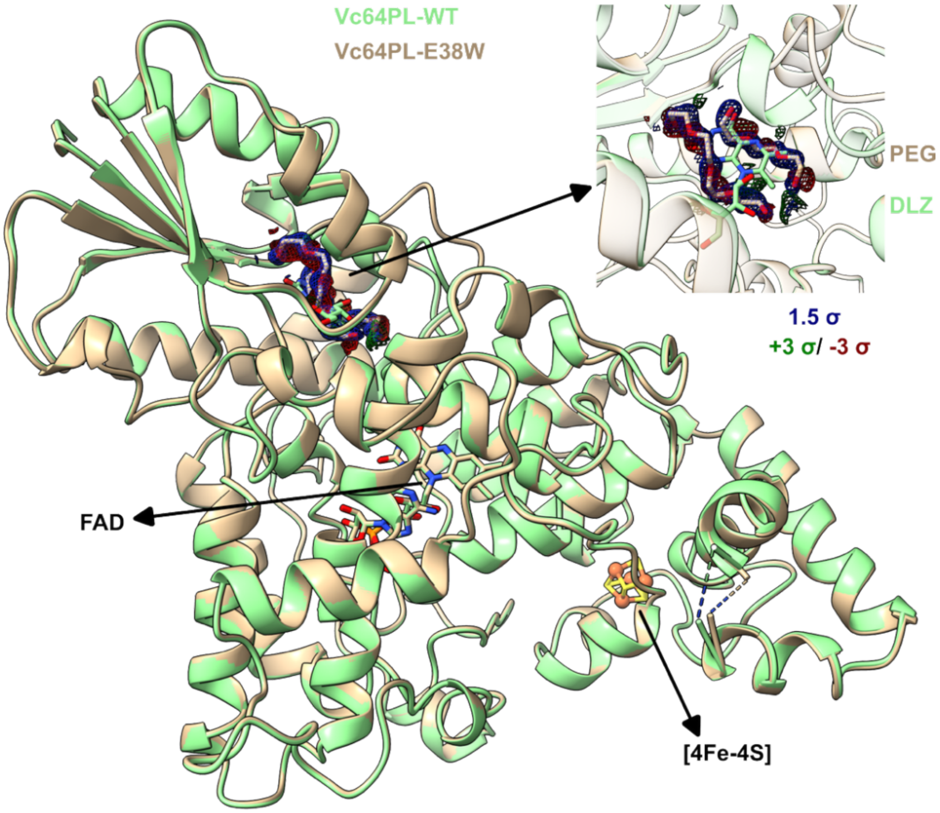


**Fig. S6 | The mutation E38W causes Vc64PL to lose DLZ binding affinity.** Shown are the cryogenic structures of Vc64PL WT and E38W (deposited under PDB codes of 32SG and 31NN, respectively) determined from disk-shaped crystals grown under the same crystallization conditions. The overall protein fold, the FAD cofactor and the [4Fe-4S] cluster are conserved. The DLZ-binding pocket of E38W is occupied by density assigned to two PEG segments (blue: 2mFo-DFc; green/red: mFo-DFc). Note that no PEG was found in the DLZ-binding pocket of rod-shaped crystals.

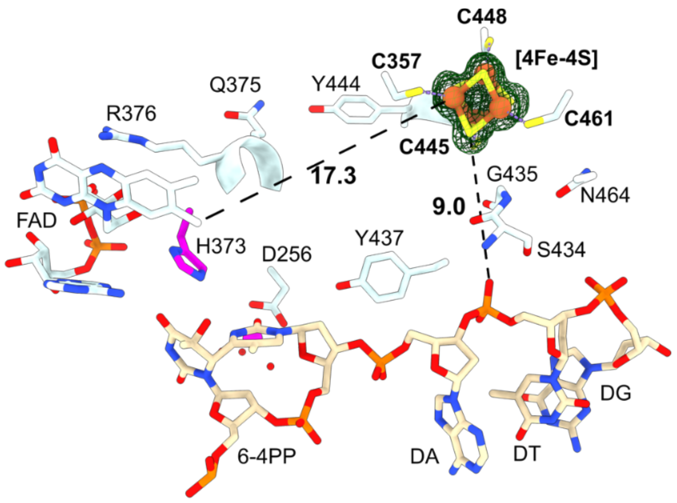


**Fig. S7 | Shortest distance from [4Fe-4S] to DNA.** Shown are residues between the [4Fe-4S] cluster and FAD or bound DNA. The distances are the minimum (atom to atom) distances from [4Fe-4S] to FAD or bound DNA in Å. The map (green, 6 σ) shown is polder map calculated with [4Fe-4S] omitted. (PDB: 31JN)

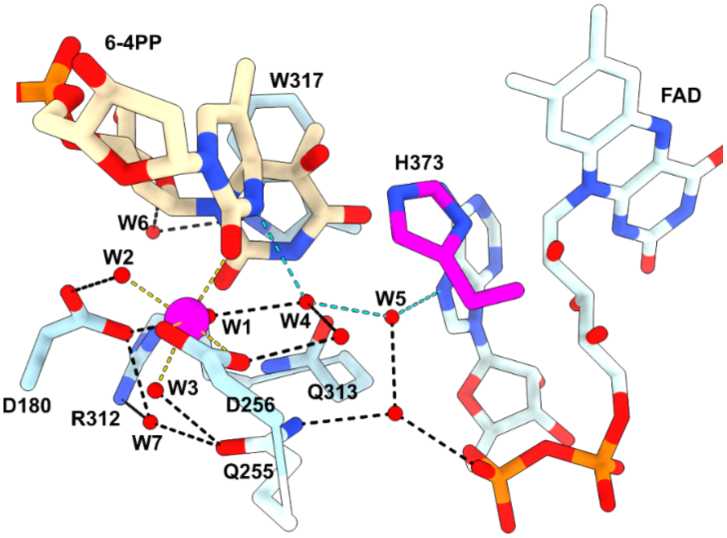


**Fig. S8 | Active-site hydrogen-bonding network.** Structural waters connect active-site residues (W317, R312, D180, Q255, D256) to form a network supporting 6-4PP. Black and cyan dashed lines represent hydrogen-bonding interactions. Gold dashed lines represent the contacts within the $Mg^{2+}$-centered octahedron shell. (PDB: 31JN)

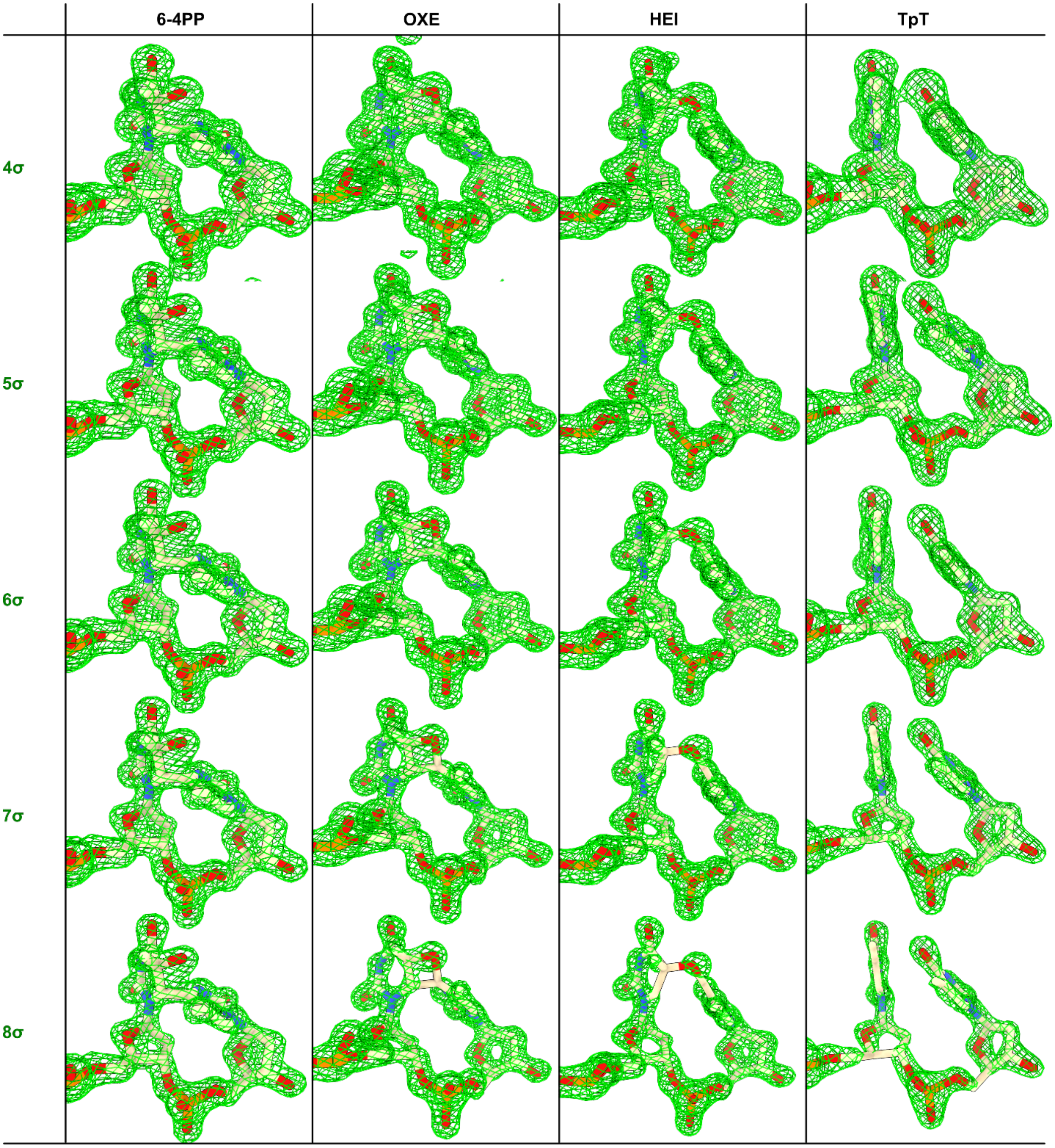


**Fig. S9 | Polder maps supporting assignments of 6-4PP, OXE, HEI and TpT.** The polder maps were calculated using (deposited) reflection data from SFX and continuous-irradiation experiments against the corresponding deposited structures (PDB entries: 31JN, 31TN, 31UD and 31IF) with the lesion, intermediate or product omitted.

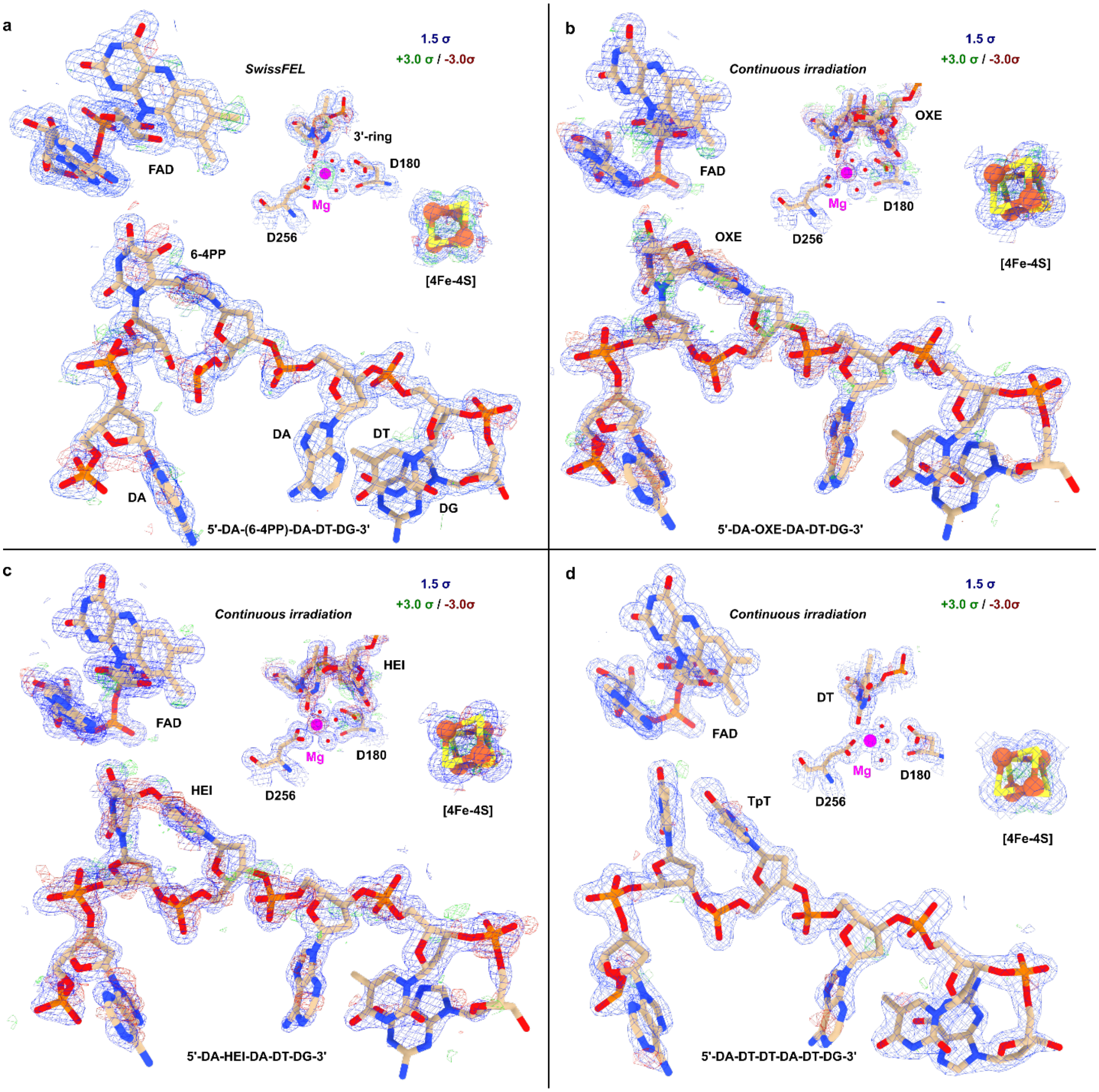


**Fig. S10 | Electron-density support for refinement.** Supplementary electron-density maps, specifically the 2mFo-DFc (light blue, 1.5 σ) and mFo-DFc maps (green 3 σ, red -3 σ), for **a,** the SFX structure (PDB: 31JN) containing 6-4PP, **b,** the structure (PDB: 31TN) containing OXE from the continuous-irradiation experiment, **c,** the structure (PDB: 31UD) containing HEI from the continuous-irradiation experiment. **d,** the structure (PDB: 31IF) containing TpT from the continuous-irradiation experiment.

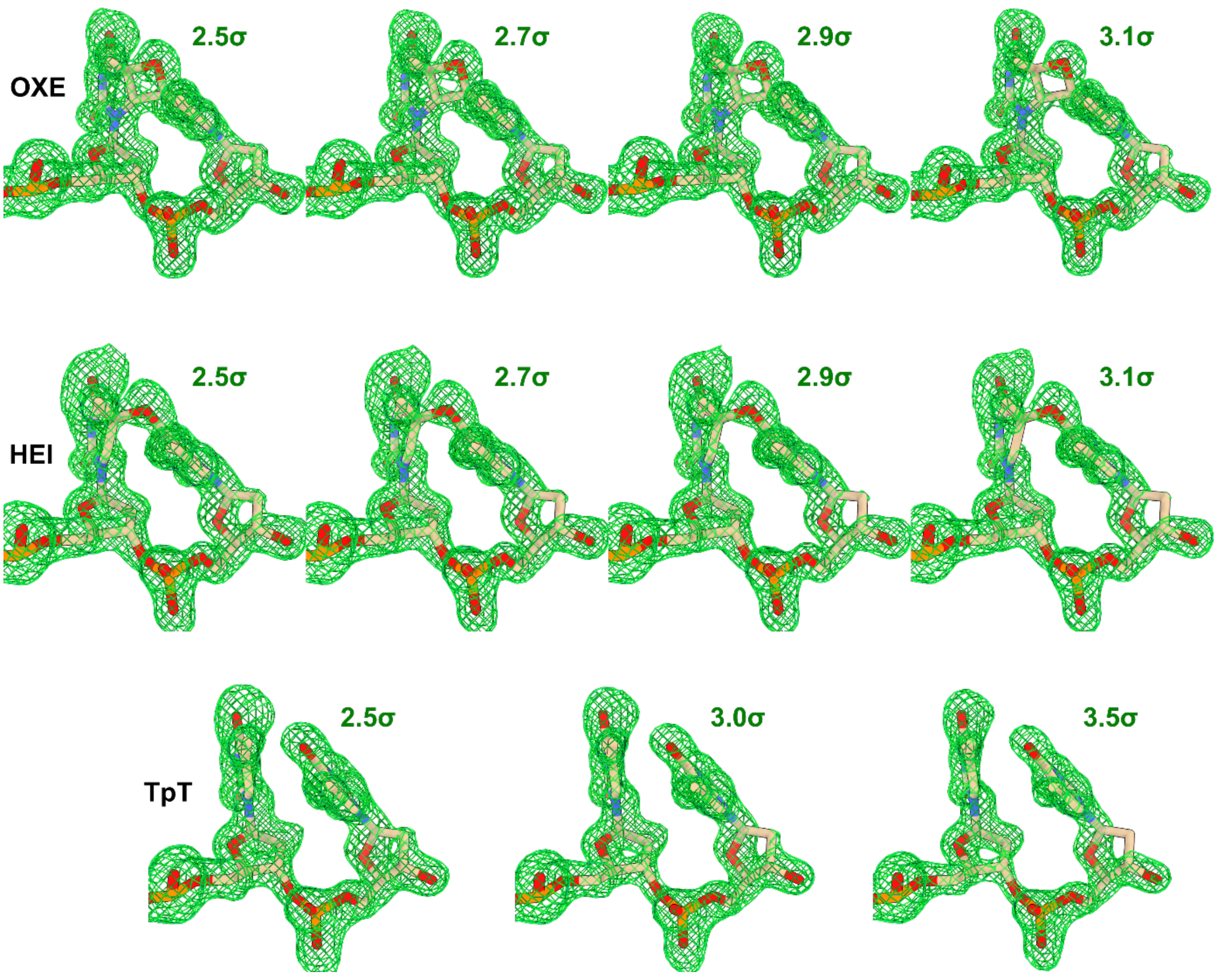


**Fig. S11 | Supplementary polder maps supporting assignment of OXE, HEI and TpT in the pre-exposure experiment.** The polder maps were calculated using reflection data from the pre-exposure experiment against deposited structures (PDB entries: 32AM, 32AN and 32AO) with the intermediate or product omitted.

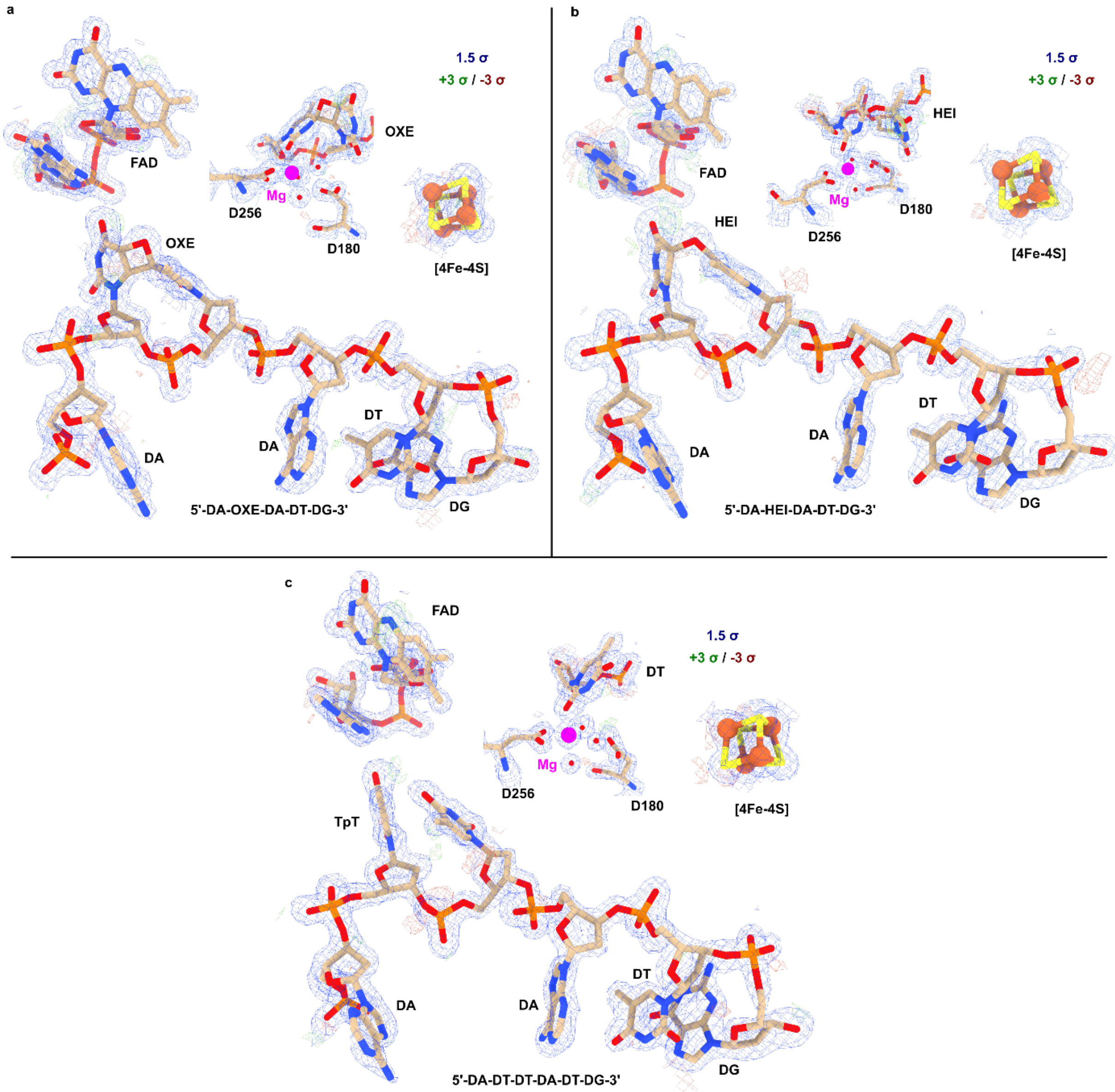


**Fig. S12 | Electron-density support for refinement of structures from the pre-exposure experiment.** Supplementary electron-density maps, specifically the 2mFo-DFc (light blue, 1.5 σ) and mFo-DFc (green 3 σ, red -3 σ) maps, for **a,** the structure (PDB: 32AM) containing OXE, **b,** the structure (PDB: 32AN) containing HEI, and **c,** the structure (PDB: 32AO) containing TpT, all from the pre-exposure experiment.

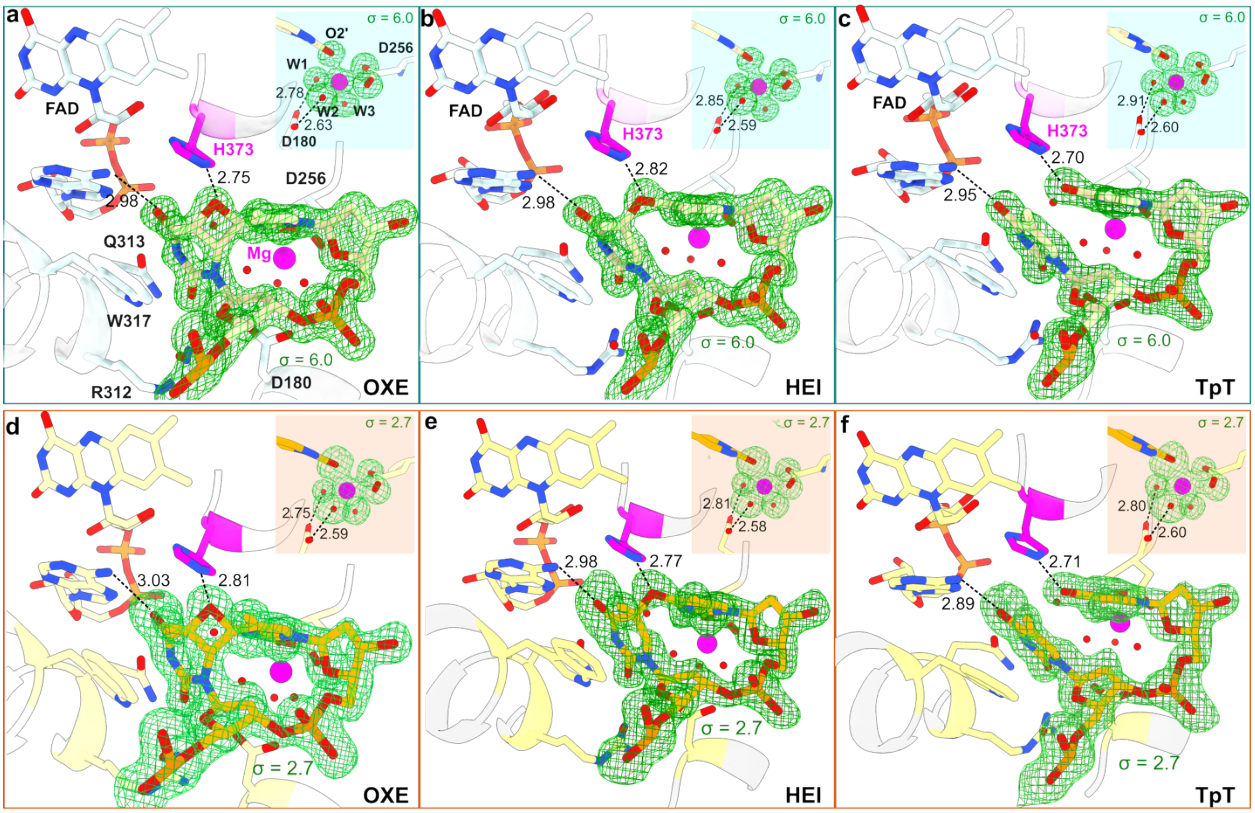


**Fig. S13 | The same sequence of intermediates is captured in independent experiments. (a-c)** The active sites of OXE, HEI and TpT from the continuous-irradiation experiment (PDBs: 31TN, 31UD, 31IF). **(d-f)** Th active sites of OXE, HEI and TpT from the pre-exposure experiment (PDBs: 32AM, 32AN, 32AO). Maps (green) shown are polder maps. (a-c) were adapted from Fig. 3 for comparison. Dashed lines represent selected hydrogen-bonding interactions. Note that other active-site hydrogen-bonding interactions shown in Fig. 1c and Fig. 3 are preserved in these structures but not shown.

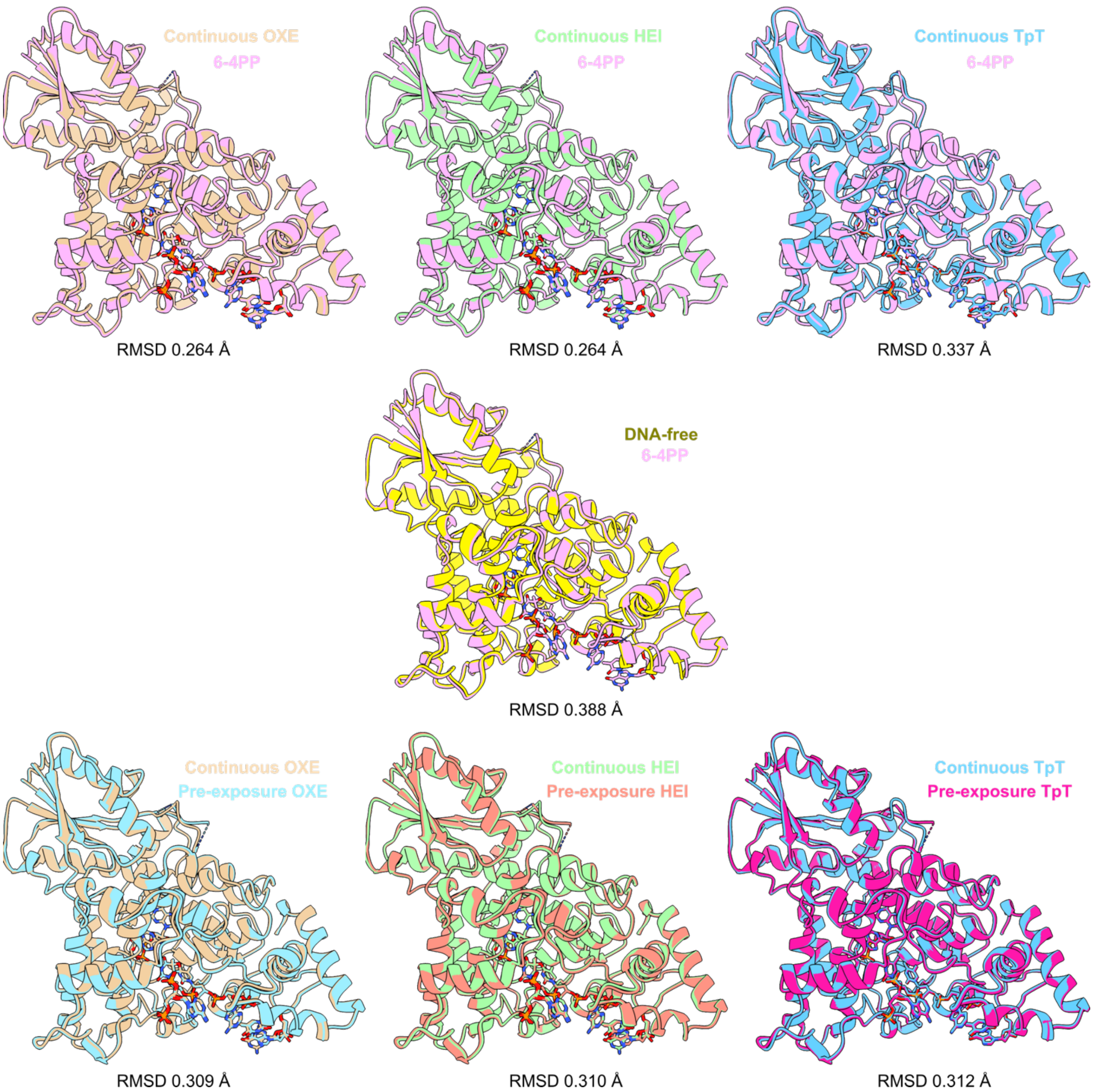


**Fig. S14 | Conservation of protein structure across different experimental conditions.** All Vc64PL-E38W structures are aligned to show that structural integrity was maintained. "6-4PP" = SFX structure containing 6-4PP (PDB: 31JN); "DNA-free" = Cryogenic E38W structure without DNA (PDB: 31NN); "Continuous" = Continuous irradiation experiment (PDBs: 31TN, 31UD, 31IF); "Pre-exposure" = Pre-exposure experiment (PDBs: 31AM, 32AN, 32AO). RMSD values for Cα alignments are listed under each comparison.

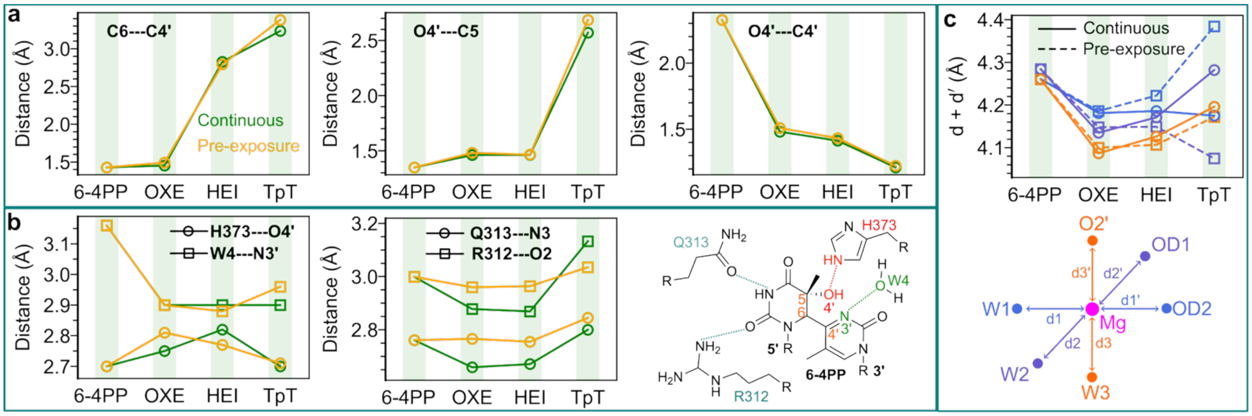


**Fig. S15 | Consistent active-site reorganization during 6-4PP repair. a,** Oxetane bond lengths. **b**, Distances between H373 and O4′, between W4 and 3′ ring N3′ and from R312/Q313 to the 5′ ring. **c**, Asymmetric contraction of the $Mg^{2+}$-coordinated octahedron during 6-4PP repair. "Continuous" = continuous-irradiation experiment; "Pre-exposure" = pre-exposure experiment. The SFX structure (PDB: 31JN) containing 6-4PP was used as the initial state.

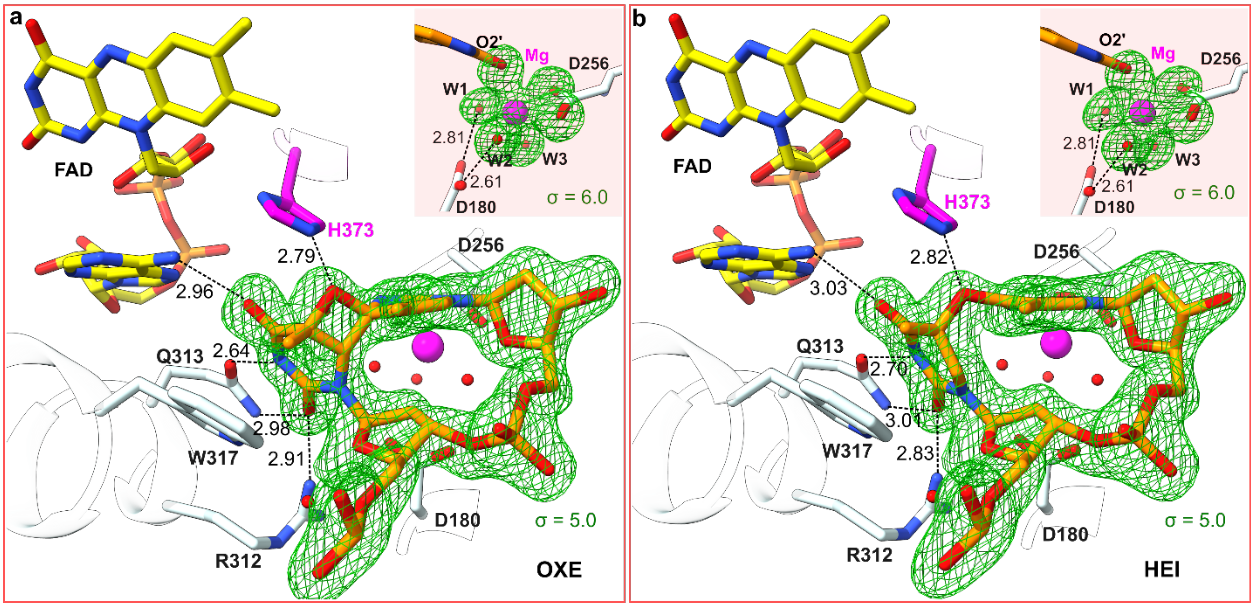


**Fig. S16 | Active-site structures of Vc64PL containing OXE and HEI. a,** active-site structure of Vc64PL with OXE. **b,** active-site structure of Vc64PL with HEI. These structures were deposited under the PDB codes: 32UC and 32UD. Major interactions between the intermediate and active-site residues were retained in both structures. The maps shown are polder maps (green). The data were collected from one disk-shaped crystal subjected to a flux of $8.8 \times 10^{12}$ photons/second with an oscillation of 0.1° and integration time of 7.5 millisecond per frame. The estimated ADW doses were 0.282 and 0.775 MGy for OXE and HEI, respectively.

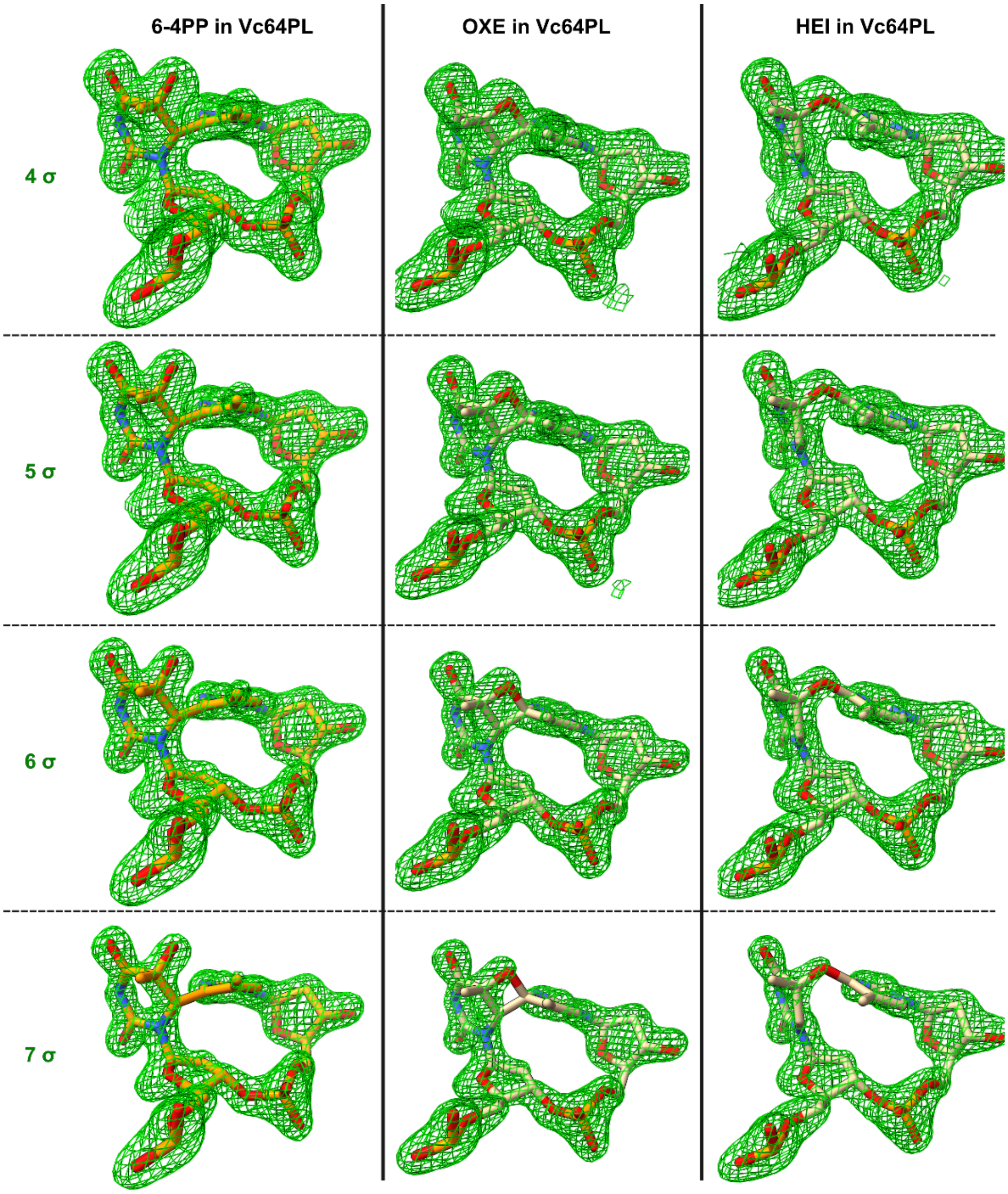


**Fig. S17 | Polder maps supporting assignments of 6-4PP, OXE and HEI in Vc64PL.** The corresponding structures were deposited under the PDB codes: 31JD, 32UC and 32UD, respectively.

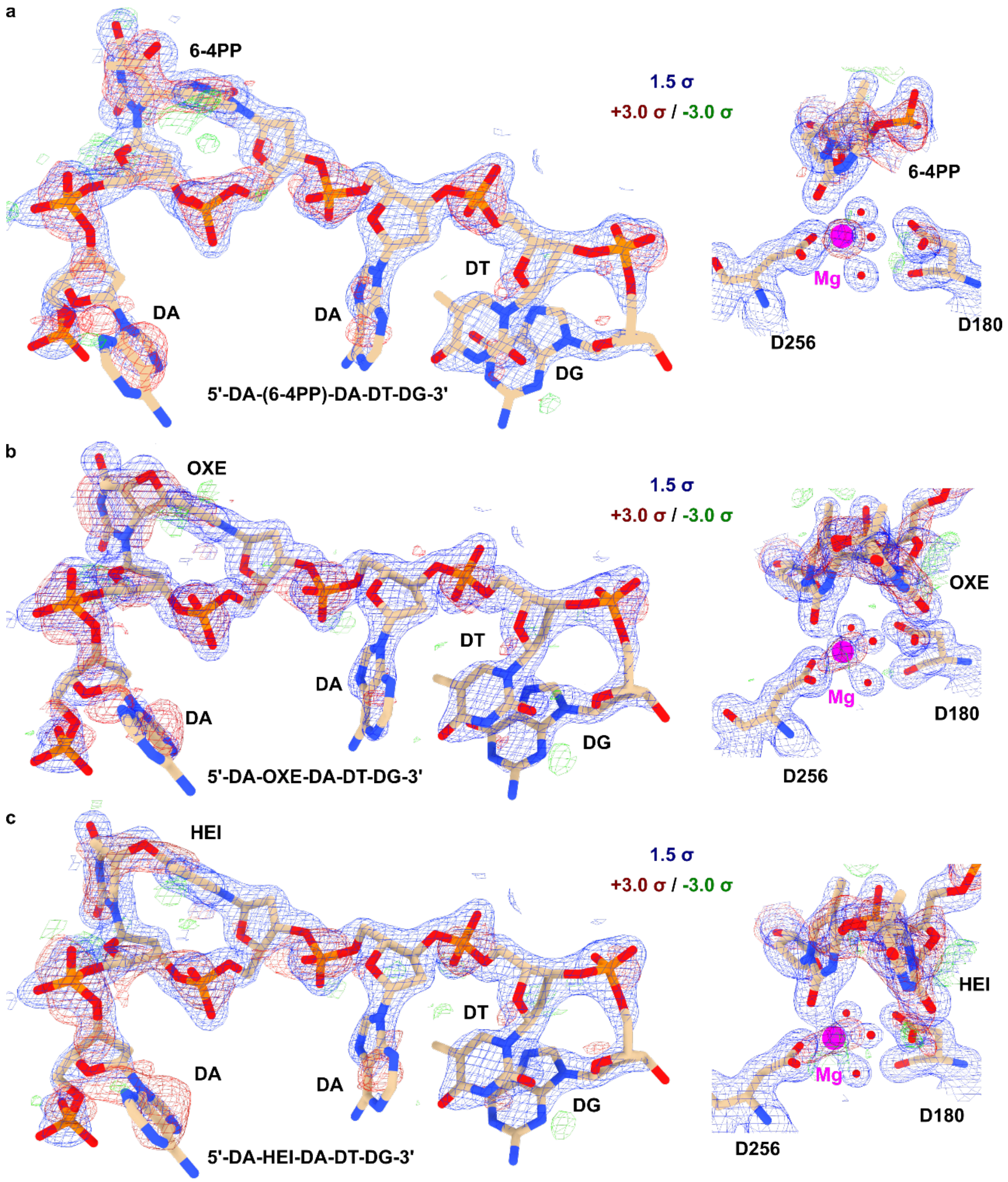


**Fig. S18 | Electron-density support for refinement of Vc64PL complex structures.** The 2mFo-DFc (light blue, 1.5 σ) and mFo-DFc (green 3 σ, red -3 σ) maps for **a,** the structure (PDB: 31JD) refined with 6-4PP, **b,** the structure (PDB: 32UC) refined with OXE, and **c,** the structure (PDB: 32UD) refined with HEI.

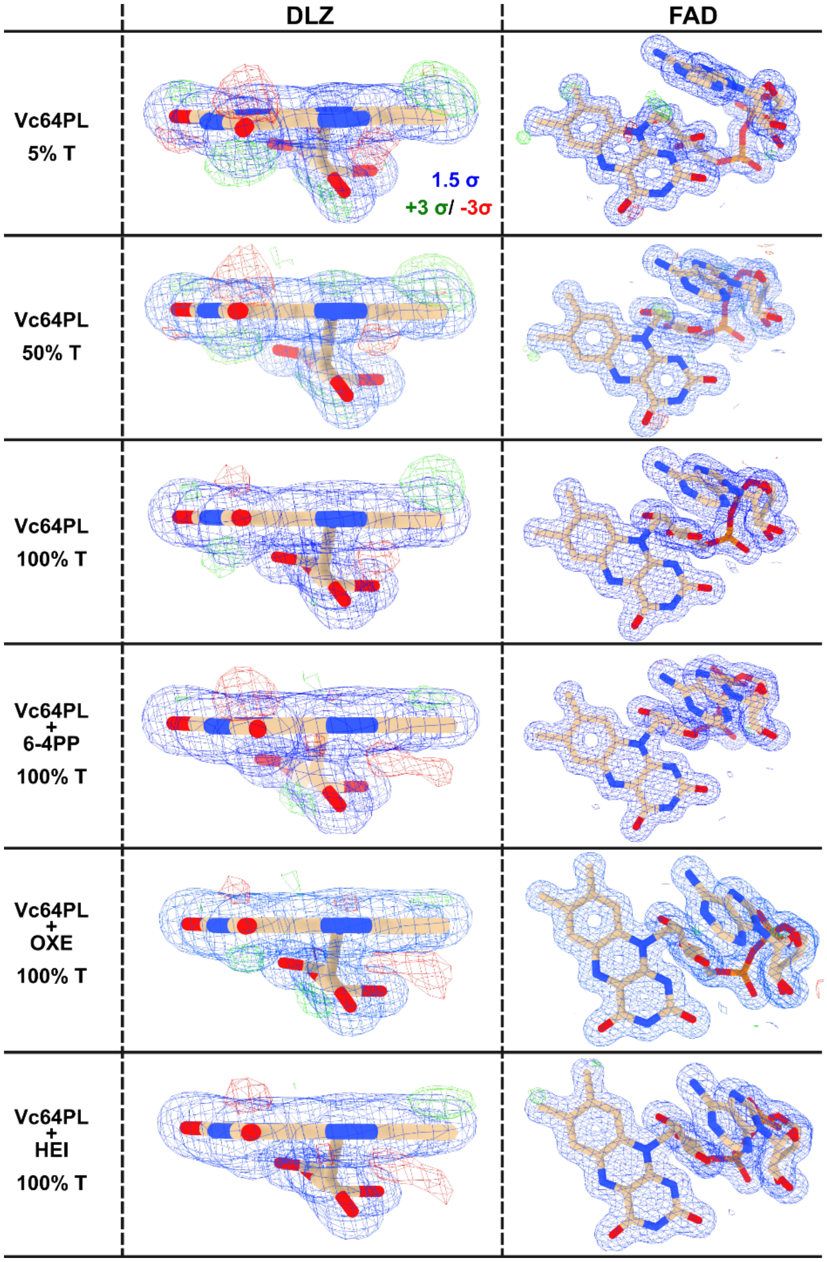


**Fig. S19 | Electron-density support for DLZ.** The maps shown are 2mFo-DFc maps (light blue, 1.5 σ) and mFo-DFc maps (green 3 σ, red -3 σ) from different Vc64PL datasets (DNA-free, 6-4PP, OXE and HEI). T indicates the X-ray transmission. 100% T corresponds to a flux of $8.8 \times 10^{12}$ photons/second. The DLZ and FAD were refined using the default restraints from Phenix monomer library. The difference maps indicate slight bending of the DLZ conjugated rings, probably due to reduction or another chemical modification. By contrast, the difference maps for FAD show negligible difference density.

**Table S1 | Data collection and refinement statistics for the serial femtosecond crystallography experiment**

| | |
|---|---|
| PDB code | **31JN** |
| Description | E38W+6-4PP |
| **Data collection and processing** | |
| Beamline | Alvra endstation, Aramis beamline (SwissFEL) |
| Detector | Jungfrau 16M |
| Wavelength (Å) | 1.03 (11.985 keV) |
| Temperature (K) | 296 |
| Space group | $P6_422$ |
| Unit-cell (a, b, c, α, β, γ) | 202.64 Å, 202.64 Å, 77.82 Å<br>90°, 90°, 120° |
| Resolution (Å) | 31.52 – 1.67 (1.71 – 1.67) |
| Number of crystals merged | 232779 |
| Total observed reflections | 194270345 (4590122) |
| Unique reflections | 108393 (10677) |
| Multiplicity | 1792.3 (430) |
| Completeness (%) | 100.00 (100.00) |
| Mean I/σ(I) | 9.87 (1.50) |
| CC1/2 | 0.998 (0.509) |
| Rsplit | 0.090 (0.774) |
| Wilson B-factor ($Å^2$) | 15.60 |
| **Refinement** | |
| Resolution (Å) | 31.52 – 1.67 (1.71 – 1.67) |
| Reflections (work+test, test) | 108123 (7580), 1997 (141) |
| $R_{work}$, $R_{free}$ | 0.1559 (0.2601), 0.1757 (0.2663) |
| No. atoms | 4872 |
| RMSD bond lengths (Å) | 0.006 |
| RMSD bond angles (°) | 0.83 |
| Ramachandran favored (%) | 97.64 |
| Ramachandran outliers (%) | 0.00 |
| Rotamer outliers (%) | 0.22 |
| Clashscore | 1.34 |
| Mean B-factor ($Å^2$) | 21.81 |

**Table S2 | Data collection and refinement statistics for the continuous-irradiation experiment.** Note that 31IF was determined from a rod-shaped crystal in our early experiment and later merged into the continuous-irradiation experimental results because it resembles a pure TpT state. We resolved an independent structure (31TQ) containing TpT from a disk-shaped crystal in the follow-up experiments. Occupancy refinement shows that 31TQ contains 61% TpT and 39% HEI.

| PDB code | **31NN** | **31TN** | **31UD** | **31TQ** | **31IF** |
|---|---|---|---|---|---|
| Description | E38W | E38W+OXE | E38W+HEI | E38W+TpT | E38W+TpT |
| **Data collection and processing** | | | | | |
| X-ray source | P14 (DESY) | | | | |
| Detector | EigerX 16M | | | | |
| Beam shape | Top-hat | | | | |
| Flux (photons/second) | $1.7 \times 10^{12}$ | $8.8 \times 10^{11}$ | | $3.5 \times 10^{12}$ | $8.8 \times 10^{12}$ |
| Integration time | 0.0075s per image | | | | |
| Oscillation | 0.1° per image | | | | |
| Wavelength (Å) | 0.976 (12.8 keV) | | | | |
| Temperature (K) | 100 | | | | |
| Space group | $P6_422$ | | | | $P2_12_12_1$ |
| Cell dimension | | | | | |
| a, b, c (Å) | 199.57, 199.57, 76.52 | 199.77, 199.77, 76.77, | 199.77, 199.77, 76.77 | 199.89, 199.89, 76.83 | 48.75, 84.18, 123.74 |
| α, β, γ (°) | 90, 90, 120 | 90, 90, 120 | 90, 90, 120 | 90, 90, 120 | 90, 90, 90 |
| Resolution (Å) | 99.78 – 1.42 (1.44 – 1.42) | 99.89 – 1.40 (1.42 – 1.40) | 99.89 – 1.40 (1.42 – 1.40) | 70.22 – 1.40 (1.42 – 1.40) | 84.14 – 1.51 (1.54 – 1.51) |
| Total reflections | 6508663 (214970) | 1575684 (77825) | 1968674 (97264) | 1573761 (77811) | 1038657 (33876) |
| Unique reflections | 166999 (8121) | 172857 (8664) | 174617 (8678) | 174906 (8688) | 79401 (3274) |
| Multiplicity | 39.0 (26.5) | 9.1 (9.0) | 11.3 (11.2) | 9.0 (9.0) | 13.1 (10.3) |
| Completeness (%) | 99.88 (98.03) | 98.61 (99.84) | 99.62 (100.00) | 99.60 (99.95) | 98.37 (82.32) |
| Mean I/σ(I) | 19.9 (1.1) | 13.5 (1.1) | 13.7 (1.1) | 13.1 (0.6) | 14.3 (0.60) |
| CC1/2 | 1.000 (0.642) | 0.999 (0.601) | 0.999 (0.593) | 0.999 (0.500) | 0.999 (0.657) |
| Rmerge | 0.114 (1.766) | 0.096 (1.175) | 0.100 (1.271) | 0.083 (1.520) | 0.072 (1.163) |
| Wilson B-factor ($Å^2$) | 17.01 | 14.77 | 15.00 | 17.36 | 22.82 |
| **Refinement** | | | | | |
| Resolution (Å) | 37.71 – 1.42 | 39.69 – 1.40 | 39.69 – 1.40 | 37.77 – 1.40 | 38.29 – 1.51 |
| Reflections (work+test, test) | 166833, 2044 | 172752, 1977 | 174515, 2007 | 174462, 2012 | 79022, 1965 |
| $R_{work}$, $R_{free}$ | 0.1665, 0.1827 | 0.1704, 0.1813 | 0.1681, 0.1781 | 0.1706, 0.1862 | 0.1980, 0.2227 |
| No. atoms | 5147 | 5157 | 5179 | 5151 | 4572 |
| RMSD bond lengths (Å) | 0.005 | 0.006 | 0.006 | 0.006 | 0.006 |
| RMSD bond angles (°) | 0.83 | 0.92 | 0.84 | 0.87 | 0.84 |
| Ramachandran favored (%) | 97.24 | 97.28 | 97.67 | 97.67 | 97.47 |
| Ramachandran outliers (%) | 0.00 | 0.00 | 0.00 | 0.00 | 0.19 |
| Rotamer outliers (%) | 0.66 | 0.44 | 0.43 | 0.43 | 0.23 |
| Clashscore | 2.03 | 2.12 | 1.99 | 2.43 | 1.29 |
| Mean B-factor ($Å^2$) | 23.84 | 20.03 | 21.25 | 23.21 | 30.05 |

**Table S3 | Data collection and refinement statistics for the pre-exposure experiment**

| PDB code | **32AM** | **32AN** | **32AO** |
|---|---|---|---|
| Description | E38W+OXE | E38W+HEI | E38W+TpT |
| Pre-exposure | No pre-exposure | 0.05 s | 1 s |
| **Data collection and processing** | | | |
| X-ray source | MicroMAX (MAX IV) | | |
| Detector | Jungfrau 9M | | |
| Beam shape | Top-hat | | |
| Flux (photons/second) | $1.3 \times 10^{14}$ | | |
| Integration time | 0.0005s per image | | |
| Oscillation | 0.1° per image | | |
| Wavelength (Å) | 0.905 (13.7 keV) | | |
| Temperature (K) | 100 | | |
| Space group | $P2_12_12_1$ | | |
| Cell dimension | | | |
| a, b, c (Å) | 50.33, 85.57, 124.96 | 50.26, 85.84, 124.54 | 50.42, 85.72, 125.07 |
| α, β, γ (°) | 90, 90, 90 | 90, 90, 90 | 90, 90, 90 |
| Resolution (Å) | 124.96 – 1.50 (1.53 – 1.50) | 124.54 – 1.50 (1.53 – 1.50) | 125.07 – 1.50 (1.53 – 1.50) |
| Total reflections | 474048 (16975) | 496086 (18035) | 472506 (16966) |
| Unique reflections | 85612 (4091) | 85142 (4154) | 86020 (4114) |
| Multiplicity | 5.5 (4.1) | 5.8 (4.3) | 5.5 (4.1) |
| Completeness (%) | 98.23 (94.83) | 97.83 (95.74) | 98.25 (94.73) |
| Mean I/σ(I) | 9.9 (0.8) | 7.3 (0.4) | 6.5 (0.5) |
| CC1/2 | 0.982 (0.577) | 0.972 (0.620) | 0.965 (0.612) |
| Rmerge | 0.096 (0.795) | 0.115 (1.183) | 0.128 (1.074) |
| Wilson B-factor (Å$^2$) | 16.77 | 16.53 | 16.66 |
| **Refinement** | | | |
| Resolution (Å) | 15.83 – 1.50 | 15.79 – 1.50 | 15.80 – 1.50 |
| Reflections (work+test, test) | 84646, 1936 | 84247, 1979 | 84893, 1969 |
| $R_{work}$, $R_{free}$ | 0.2204, 0.2408 | 0.2285, 0.2626 | 0.2425, 0.2593 |
| No. atoms | 4734 | 4712 | 4641 |
| RMSD bond lengths (Å) | 0.007 | 0.007 | 0.006 |
| RMSD bond angles (°) | 0.95 | 0.97 | 0.85 |
| Ramachandran favored (%) | 97.83 | 97.83 | 97.64 |
| Ramachandran outliers (%) | 0.00 | 0.00 | 0.00 |
| Rotamer outliers (%) | 0.69 | 0.46 | 0.93 |
| Clashscore | 2.25 | 2.01 | 2.25 |
| Mean B-factor (Å$^2$) | 20.37 | 21.14 | 22.93 |

**Table S4 | Data collection and refinement statistics for Vc64PL structures.** The structure of wild-type Vc64PL has been reported previously[4,13]. Here we resolved a new structure (32SG) with higher resolution. The Vc64PL:6-4PP crystal was intentionally exposed to the full flux of the P14 X-ray beam to verify whether 6-4PP was repaired as efficiently as that in E38W (31IF). The calculated ADW doses for 31JD, 32UC and 32UD were 0.141, 0.282 and 0.775 MGy, respectively.

| PDB code | **32SG** | **31JD** | **32UC** | **32UD** |
|---|---|---|---|---|
| Description | Vc64PL | Vc64PL+6-4PP | Vc64PL+OXE | Vc64PL+HEI |
| **Data collection and processing** | | | | |
| X-ray source | P14 (DESY) | | | |
| Detector | EigerX 16M | | | |
| Beam shape | Top-hat | | | |
| Flux (photons/second) | $4.4 \times 10^{12}$ | $8.8 \times 10^{12}$ | | |
| Exposure time | 0.0075 s/image | | | |
| Oscillation | 0.1 degree/image | | | |
| Wavelength (Å) | 0.976 (12.8 keV) | | | |
| Temperature (K) | 100 | | | |
| Space group | $P6_422$ | | | |
| a, b, c (Å) | 200.01, 200.01, 77.08 | 202.26, 202.26, 77.40 | 202.26, 202.26, 77.40 | 202.26, 202.26, 77.40 |
| α, β, γ (°) | 90, 90, 120 | 90, 90, 120 | 90, 90, 120 | 90, 90, 120 |
| Resolution (Å) | 100.03 – 1.38 (1.40 – 1.38) | 101.13 – 1.52 (1.55 – 1.52) | 61.47 – 1.55 (1.58 – 1.55) | 101.13 – 1.60 (1.63 – 1.60) |
| Total reflections | 7394221 (324928) | 959028 (47559) | 903120 (44380) | 1233825 (60468) |
| Unique reflections | 184186 (9097) | 140522 (7005) | 129842 (6389) | 122003 (6011) |
| Multiplicity | 40.1 (35.7) | 6.8 (6.8) | 7.0 (6.9) | 10.1 (10.1) |
| Completeness (%) | 99.99 (99.96) | 98.99 (99.83) | 96.92 (96.67) | 100.00 (100.00) |
| Mean I/σ(I) | 17.0 (0.6) | 15.9 (0.6) | 14.2 (0.5) | 16.7 (0.5) |
| CC1/2 | 1.000 (0.544) | 0.999 (0.619) | 0.999 (0.529) | 0.999 (0.539) |
| Rmerge | 0.100 (1.959) | 0.055 (1.081) | 0.060 (1.140) | 0.063 (1.682) |
| Wilson B-factor ($Å^2$) | 17.43 | 24.27 | 25.28 | 28.22 |
| **Refinement** | | | | |
| Resolution (Å) | 39.75 – 1.38 | 42.33 – 1.52 | 48.58 – 1.55 | 70.80 – 1.60 |
| Reflections (work+test, test) | 183692, 1995 | 140277, 1988 | 129183, 1944 | 121779, 1987 |
| $R_{work}$, $R_{free}$ | 0.1664, 0.1763 | 0.1763, 0.1844 | 0.1731, 0.1852 | 0.1739. 0.1903 |
| No. atoms | 5005 | 5017 | 5012 | 5006 |
| RMSD bond lengths (Å) | 0.006 | 0.006 | 0.006 | 0.006 |
| RMSD bond angles (°) | 0.88 | 0.87 | 0.88 | 0.89 |
| Ramachandran favored (%) | 97.63 | 97.47 | 97.28 | 97.28 |
| Ramachandran outliers (%) | 0.00 | 0.00 | 0.00 | 0.00 |
| Rotamer outliers (%) | 0.22 | 0.22 | 0.22 | 0.22 |
| Clashscore | 1.02 | 2.98 | 2.54 | 2.44 |
| Mean B-factor ($Å^2$) | 23.41 | 30.64 | 30.32 | 33.47 |

**Table S5 | Results of occupancy refinement from the continuous-irradiation experiment.** Dose calculated using RADDOSE-3D[14] is reported as the average diffraction weighted (AWD) dose. The total exposure time for each crystal was 27 s (3600 images, 7.5 ms per image). The last data point (0.7167 MGy) contained 1200 images (images 2400-3600) to ensure completeness because the data was collected from a rod-shaped crystal (Fig. S1a). All other data wedges, collected from disk-shaped crystals, contained 800 images, so the cumulative exposure time for each data point ranged from 6 to 27 s. Under the adopted experimental conditions, the intermediate occupancies showed a strong dependency on the X-ray dose rather than on exposure time. Note that the zeros in this table mean that the state was not included in the corresponding occupancy refinement, not that its occupancy refined to zero.

| ADW Dose (MGy) | 6-4PP | OXE | HEI | TpT |
|---|---|---|---|---|
| 0.0019 | 0.60 | 0.40 | 0 | 0 |
| 0.0028 | 0.56 | 0.44 | 0 | 0 |
| 0.0075 | 0 | 0.73 | 0.27 | 0 |
| 0.0188 | 0 | 0.74 | 0.26 | 0 |
| 0.0282 | 0 | 0.54 | 0.46 | 0 |
| 0.0376 | 0 | 0.50 | 0.50 | 0 |
| 0.0469 | 0 | 0.36 | 0.64 | 0 |
| 0.0563 | 0 | 0.28 | 0.73 | 0 |
| 0.0657 | 0 | 0.23 | 0.77 | 0 |
| 0.0751 | 0 | 0.16 | 0.84 | 0 |
| 0.0845 | 0 | 0.11 | 0.78 | 0.11 |
| 0.0939 | 0 | 0 | 0.65 | 0.35 |
| 0.1127 | 0 | 0 | 0.68 | 0.32 |
| 0.1314 | 0 | 0 | 0.66 | 0.34 |
| 0.1408 | 0 | 0 | 0.66 | 0.34 |
| 0.1502 | 0 | 0 | 0.57 | 0.43 |
| 0.1690 | 0 | 0 | 0.62 | 0.38 |
| 0.1878 | 0 | 0 | 0.47 | 0.53 |
| 0.1971 | 0 | 0 | 0.62 | 0.38 |
| 0.2253 | 0 | 0 | 0.54 | 0.46 |
| 0.2535 | 0 | 0 | 0.57 | 0.43 |
| 0.2629 | 0 | 0 | 0.43 | 0.57 |
| 0.3004 | 0 | 0 | 0.39 | 0.61 |
| 0.3380 | 0 | 0 | 0.39 | 0.61 |
| 0.7167 | 0 | 0 | 0 | 1 |

**Table S6 | Raddose-3D simulation to evaluate rotational dependence of X-ray absorption.** The table shows values of average diffraction weighted (AWD) dose calculated by Raddose-3D when choosing different starting angular positions. A test-wedge size of 40° was chosen for better comparison with real experimental setup. A top-hat 12.8 keV X-ray beam with a flux of $8.8 \times 10^{12}$ photons/second and a diameter of 300 µm is assumed to irradiate a disk-shaped crystal with a dimension of $300 \times 300 \times 80$ $\mu m^3$ for 3 seconds as the crystal is rotated by 40°. The calculated AWD doses show that the variation of X-ray absorption due to rotation is less than 5% in our continuous-irradiation experiment.

| Starting angular position | Ending angular position | Calculated AWD dose (MGy) |
|---|---|---|
| 0° | 40° | 0.0923 |
| 40° | 80° | 0.0923 |
| 80° | 120° | 0.0937 |
| 120° | 160° | 0.0960 |
| 160° | 200° | 0.0948 |
| 200° | 240° | 0.0916 |
| 240° | 280° | 0.0933 |
| 280° | 320° | 0.0951 |
| 320° | 360° | 0.0962 |
| Average | | 0.0939MGy/40° |

RADDOSE-3D input file for calculating absorbed X-ray dose in the continuous-irradiation experiment

*Crystal*
*Type Cuboid*
*Dimensions 300 300 80*
*PixelsPerMicron 0.5*
*AbsCoefCalc RD3D*
*UnitCell 199.7 199.7 76.7 90 90 120*
*NumMonomers 12*
*NumResidues 516*
*ProteinHeavyAtoms Fe 4 Mg 1 S 33 P 2*
*NumDNA 8*
*SolventHeavyConc Mg 0.5 S 1000 Na 100 Cl 101.*
*SolventFraction 0.66*
*Beam*
*Type TopHat*
*Flux 8.8e12*
*Energy 12.8*
*Collimation Circular 300 300*
*Wedge 0 80*
*ExposureTime 6*
*AngularResolution 1*

RADDOSE-3D input file for calculating absorbed X-ray dose in pre-exposure experiment

*Crystal*
*Type Cuboid*
*Dimensions 80 200 50*
*PixelsPerMicron 0.5*
*AbsCoefCalc RD3D*
*UnitCell 50.3 85.7 125 90 90 90*
*NumMonomers 4*
*NumResidues 516*
*ProteinHeavyAtoms Fe 4 Mg 1 S 33 P 2*
*NumDNA 8*
*SolventHeavyConc Mg 5 Na 100 Cl 110*
*SolventFraction 0.45*
*Beam*
*Type TopHat*
*Flux 1.3e14*
*Energy 13.7*
*Collimation Circular 200 200*
*Wedge 0 180*
*ExposureTime 0.9*
*AngularResolution 0.1*
